# Tumor-versus-non-tumor T cell clonotype enrichment enables antigen-agnostic prioritization of tumor-reactive TCR candidates, supported by single-cell and spatial phenotyping

**DOI:** 10.64898/2026.09.18.750832

**Authors:** V. Seitz, J. Franzen, K. Gennermann, B. Hirsch, S. Schaper, A. Dröge, M. Dreger, N. Genzel, S. Plassmann, D. Bents, J. Glökler, T. Conrad, C. Loddenkemper, K. Schmelz, M. Morkel, A. Trinks, T. Krieger, D. Kainmueller, M. Hummel, S. Hennig, D. Horst, V. Lennerz, S. Elezkurtaj

**Affiliations:** Charité – Universitätsmedizin Berlin, Institute of Pathology, Berlin, Germany; HS Diagnomics GmbH, Berlin, Germany; Max-Delbrück-Center for Molecular Medicine in the Helmholtz Association and Helmholtz Imaging, Berlin, Germany; Digital Engineering Faculty, University of Potsdam, Germany; Berlin Institute of Health (BIH) at Charité – Universitätsmedizin Berlin, Berlin, Germany; Pathotres GbR, Berlin, Germany; Charité – Universitätsmedizin Berlin, Charité 3R, Primary Tissue Pipeline, Berlin, Germany

**Keywords:** Tumor-reactive T cell receptors, T cell receptor repertoire, Antigen-agnostic TCR discovery, Spatial transcriptomics, Pancreatic ductal adenocarcinoma, Tumor microenvironment, T cell clonotypes, TCR-T cell therapy

## Abstract

**Introduction:** Identifying tumor-reactive T cell receptors (TCRs) remains challenging, particularly when tumor antigens are unknown. We investigated tumor-to-non-tumor (T/N) clonotype enrichment as an antigen-agnostic strategy for prioritizing candidate tumor-reactive TCRs, supported by single-cell and spatial phenotyping.

**Methods:** We performed bulk and single-cell TCR repertoire sequencing of paired tumor and adjacent non-tumor tissues across 15 cases comprising lung, pancreatic, breast, and colorectal cancers. Selected clonotypes were further characterized by single-cell gene expression profiling and Xenium spatial transcriptomics in two pancreatic cancer cases.

**Results:** Tumor-enriched clonotypes included Cluster-A, a public HLA-A*02:01-restricted TCR cluster preferentially enriched in tumors across 14 HLA-A*02:01-positive patients, with previously demonstrated tumor recognition. In one pancreatic cancer case, spatial analysis revealed that Cluster-A CD8⁺ T cells were exclusively detected in tumor tissue, preferentially localized near malignant epithelial cells, and exhibited an antigen-experienced effector-memory phenotype with elevated granzyme A and K expression. In another pancreatic cancer case lacking Cluster-A, the six most frequent CD8⁺ T cell clonotypes showed a graded association between T/N enrichment and tumor-associated features, with higher T/N ratios correlating with closer proximity to malignant cells and increased granzyme A and K expression.

**Conclusion:** These findings support the T/N ratio as a robust primary criterion for prioritizing candidate tumor-reactive TCRs. Integration of single-cell and spatial transcriptomics provides complementary biological evidence of tumor association and a framework for selecting therapeutic TCR candidates when functional validation is limited.

## Introduction

Adoptive cell therapy (ACT) using T cells engineered with tumor-reactive T cell receptors (TCR-T cells) offers a promising therapeutic approach for solid cancers (1–3). While chimeric antigen receptors (CARs) are currently restricted to a limited number of suitable cell-surface targets, such as CD19 and CD20 on B cells and BCMA on plasma cells, TCRs can recognize a much broader range of antigens presented by HLA molecules (4, 5).This enables TCR-T cells to target a broader range of tumor-specific epitopes with superior sensitivity (6). However, to fully realize the therapeutic potential of tumor-specific TCRs, their efficient, precise, and ideally personalized identification is crucial in order to produce engineered autologous cytotoxic T cells.

There are two strategies to select tumor-specific therapeutic TCRs: antigen-directed and antigen-agnostic. Antigen-directed approaches so far target only a narrow set of antigens (e.g. PRAME, NY-ES0-1, MAGE) paired with common HLAs, primarily HLA-A*02:01 (1). Alternatively, these strategies can focus on the identification of personalized neoantigens to stimulate and expand tumor-specific T cells, clone their TCRs and introduce them into autologous T cells for ACT (7, 8). However, the difficulty to reliably identify tumor antigens that elicit potent specific and efficacious T cell responses limits its practical application.

Our antigen-agnostic approach, originally described by us in 2015, bypasses these difficulties and leverages the well-documented phenomenon that tumor-specific T cells undergo clonal expansion upon target antigen encounter and activation within the tumor microenvironment (9, 10). To identify efficacious tumor specific TCR clonotypes, we established and validated a robust and efficient process to select tumor-expanded TCRs that takes advantage of the distinct biology of T cells engaging in tumor cell recognition and killing including protein– and transcriptome level analyses (10).

In this workflow, tumor-enriched TCR clonotypes are identified based on their numerical dominance within tumor-infiltrating lymphocytes (TILs) and a high tumor-to-non-tumor (T/N) frequency ratio. Of note, the initial identification of tumor-expanded T cell clonotypes does not rely on single-cell technologies, but can also be performed using quantitatively optimized multiplex primer-based T cell profiling from bulk DNA of fresh or even FFPE-preserved tumor and adjacent non-tumor tissue (10–12). TCR profiling of PD-1⁺ T cells sorted from fresh tissue provides additional evidence of an activated/exhausted phenotype at the protein level. Subsequently, single-cell sequencing (scRNA-Seq) profiles are used as a surrogate for functional tumor reactivity to substantiate our primary selection of tumor-reactive clonotypes. Cells belonging to these clonotypes exhibit gene signatures of activation and cytotoxicity, as well as of chronic activation mirroring clonotype differentiation trajectories from activation to terminal differentiation, dysfunction and/or exhaustion (10).

Functional validation of candidate TCRs remains a major bottleneck, as it typically relies on viable tumor material and robust in vitro assay systems, which are often unavailable or difficult to establish. We therefore asked whether spatially resolved single-cell *in situ* analyses could provide complementary evidence for the tumor association of selected clonotypes. Specifically, we explored whether tumor-enriched T cells exhibit characteristic spatial and transcriptional features within the tumor microenvironment, including localization to tumor regions, proximity to malignant cells, and gene expression programs consistent with those observed by scRNA-seq.

The application of our analysis process to tumor cases across different patients and various tumor types enabled us to uncover T cell clusters with identical or near-identical TCRs that were restricted to specific HLA types (13). Notably, our lead cluster (T cell Cluster-A) demonstrated HLA-A*02:01-restricted pan-cancer reactivity (13). Such clonotypes, detected in several individuals, are referred to as common or public TCRs, in contrast to private TCRs restricted to a single individual.

Here, we extended our antigen-agnostic TCR profiling framework by integrating single-cell analyses across 15 tumor cases, including 14 cases harboring Cluster-A TCR clonotypes and PDAC_10, in which the six most abundant private clonotypes were investigated. In two selected PDAC cases, we additionally integrated Xenium spatial transcriptomics to investigate the spatial localization and transcriptional phenotype of these clonotypes within the tumor microenvironment at single-cell resolution (14, 15). Cluster-A CD8^+^ T cells localized to tumor-associated regions, were absent from matched normal pancreas, and were significantly closer to malignant epithelial cells than naïve/central memory (CM)-like CD8^+^ T cells, while remaining more distant from non-malignant ductal epithelium. Cluster-A T cells were also detected within tertiary lymphoid structures (TLS) and displayed a cytotoxic effector-memory gene signature. Although some Cluster-A TCRs have been reported to cross-react with the Epstein–Barr virus (EBV)-derived antigen LMP2A, EBER *in situ* hybridization confirmed the absence of EBV-positive cells in the analyzed tumor. Accordingly, HLA-matched tumor cell lines recognized by Cluster-A TCR-T cells were not infected by EBV, as determined by RNA-Seq (13).

This observation is consistent with the emerging concept that microbial–tumor cross-reactivity through molecular mimicry is a common feature of T cell immunity and can contribute to tumor recognition rather than preclude it (16).

Spatial analysis of the six most abundant private clonotypes in a separate PDAC case further revealed a continuous tumor-reactivity spectrum linking increasing tumor-versus-non-tumor (T/N) ratios with closer proximity to malignant cells and higher expression of GZMA and GZMK (Fig. 1).

**Figure 1:**
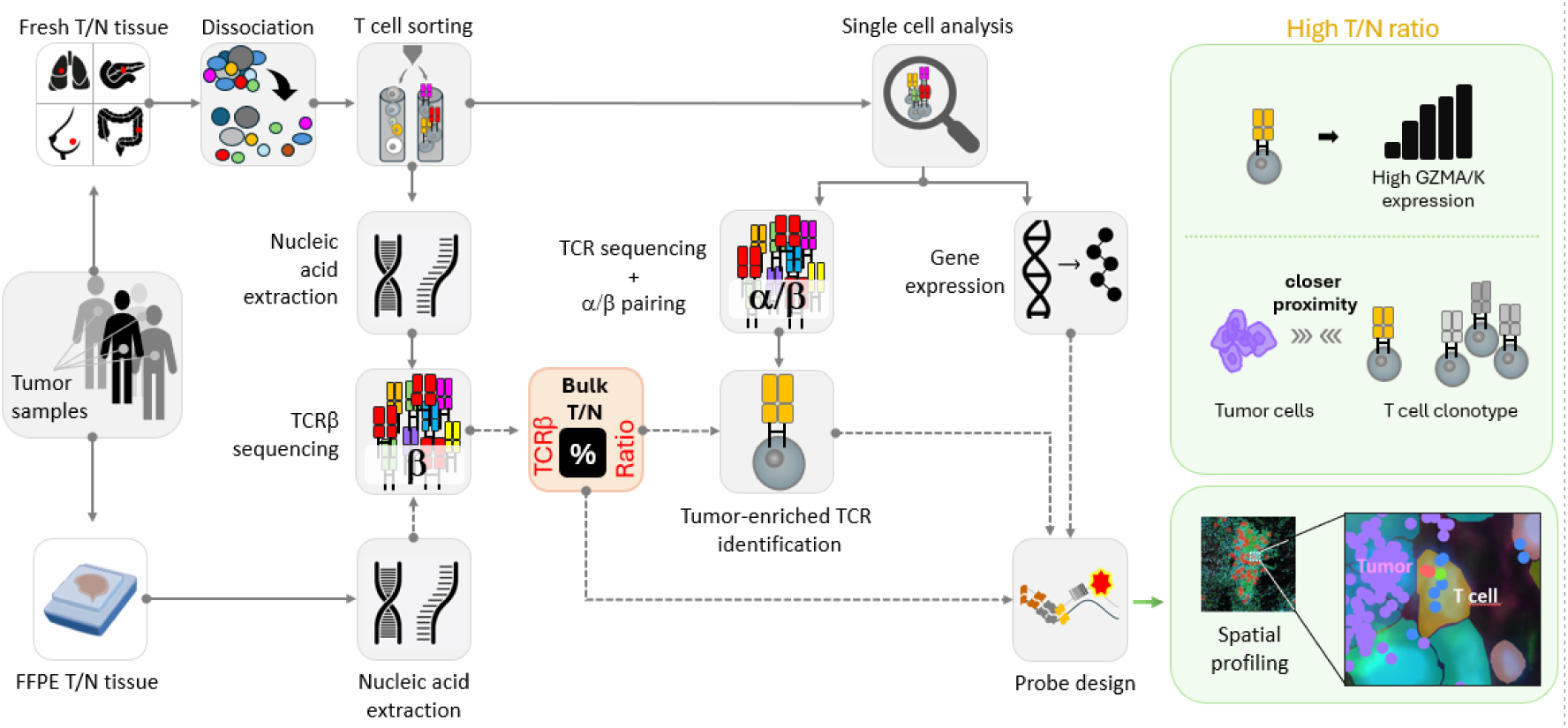
Antigen-agnostic identification and spatial validation of tumor-specific T cell candidates. Tumor-to-non-tumor (T/N) frequency ratios obtained from bulk TCR repertoire profiling were used to prioritize candidate tumor-reactive clonotypes across solid tumors. Single-cell TCR/RNA sequencing linked selected clonotypes to their transcriptional phenotype and enabled reconstruction of paired αβ TCRs. Xenium spatial transcriptomics subsequently mapped these clonotypes within the tumor microenvironment, demonstrating preferential localization in closer proximity to malignant epithelial cells than bystander T cells. In the present study, tumor-enriched clonotypes were also associated with elevated cytotoxic gene expression (GZMA/GZMK), providing an orthogonal spatial validation framework for prioritizing therapeutic TCR candidates.

Together, our findings nominate spatial profiling as an orthogonal feature that complements tumor-versus-non-tumor (T/N) repertoire enrichment for the antigen-agnostic prioritization of candidate tumor-reactive TCR clonotypes.

## Material and Methods

### Key resources table

For reproducibility, the key resources table (Table 1) lists reagents, antibodies, software, instrumentation, etc. as they are referred to in the following chapters.

**Table 1:**
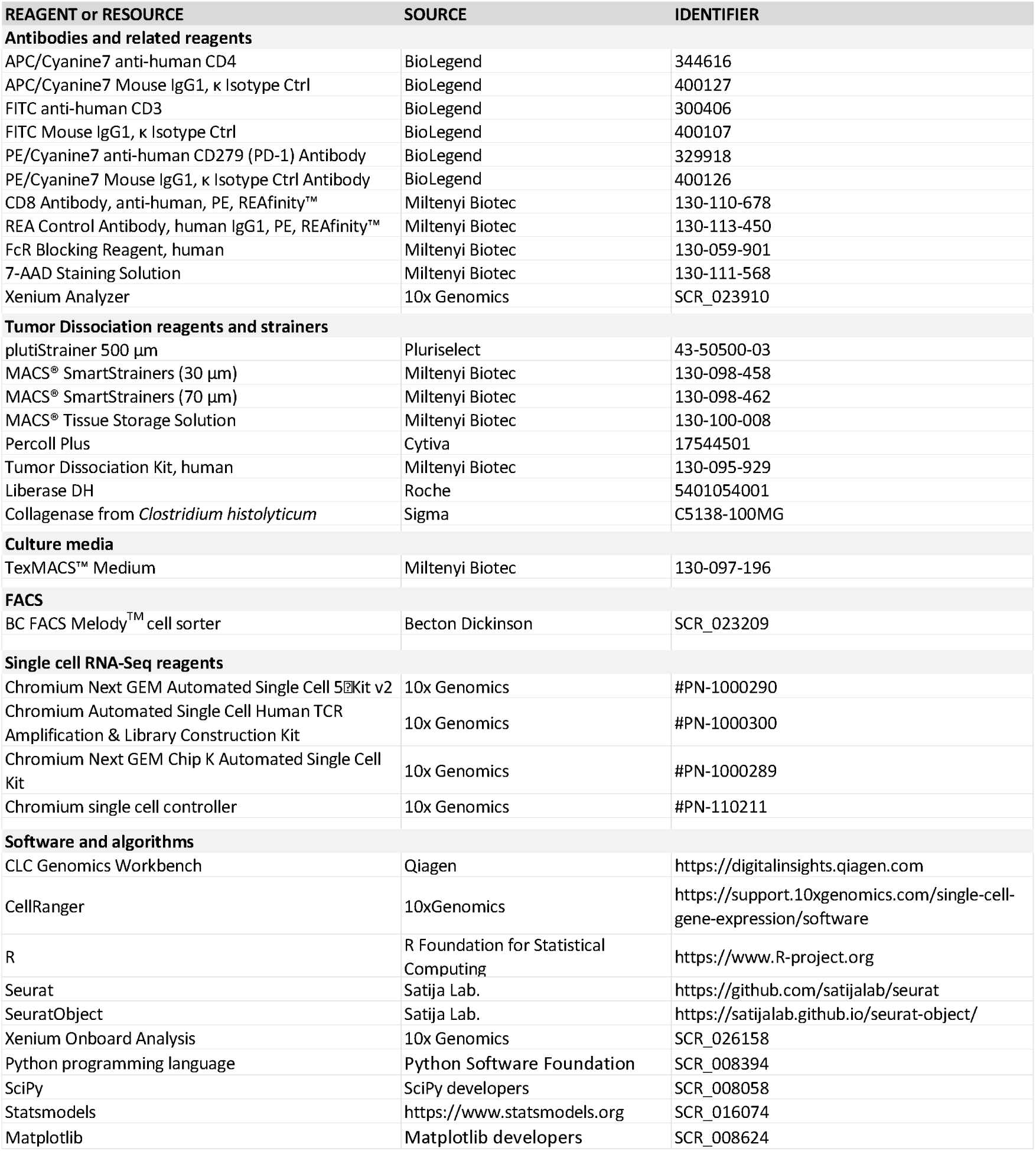
Key resources table.

| REAGENT or RESOURCE | SOURCE | IDENTIFIER |
|---|---|---|
| **Antibodies and related reagents** |  |  |
| APC/Cyanine7 anti-human CD4 | BioLegend | 344616 |
| APC/Cyanine7 Mouse IgG1, κ Isotype Ctrl | BioLegend | 400127 |
| FITC anti-human CD3 | BioLegend | 300406 |
| FITC Mouse IgG1, κ Isotype Ctrl | BioLegend | 400107 |
| PE/Cyanine7 anti-human CD279 (PD-1) Antibody | BioLegend | 329918 |
| PE/Cyanine7 Mouse IgG1, κ Isotype Ctrl Antibody | BioLegend | 400126 |
| CD8 Antibody, anti-human, PE, REAfinity™ | Miltenyi Biotec | 130-110-678 |
| REA Control Antibody, human IgG1, PE, REAfinity™ | Miltenyi Biotec | 130-113-450 |
| FcR Blocking Reagent, human | Miltenyi Biotec | 130-059-901 |
| 7-AAD Staining Solution | Miltenyi Biotec | 130-111-568 |
| Xenium Analyzer | 10x Genomics | SCR_023910 |
| **Tumor Dissociation reagents and strainers** |  |  |
| plutiStrainer 500 µm | Pluriselect | 43-50500-03 |
| MACS® SmartStrainers (30 µm) | Miltenyi Biotec | 130-098-458 |
| MACS® SmartStrainers (70 µm) | Miltenyi Biotec | 130-098-462 |
| MACS® Tissue Storage Solution | Miltenyi Biotec | 130-100-008 |
| Percoll Plus | Cytiva | 17544501 |
| Tumor Dissociation Kit, human | Miltenyi Biotec | 130-095-929 |
| Liberase DH | Roche | 5401054001 |
| Collagenase from *Clostridium histolyticum* | Sigma | C5138-100MG |
| **Culture media** |  |  |
| TexMACS™ Medium | Miltenyi Biotec | 130-097-196 |
| **FACS** |  |  |
| BC FACS Melody<sup>TM</sup> cell sorter | Becton Dickinson | SCR_023209 |
| **Single cell RNA-Seq reagents** |  |  |
| Chromium Next GEM Automated Single Cell 5�Kit v2 | 10x Genomics | #PN-1000290 |
| Chromium Automated Single Cell Human TCR Amplification & Library Construction Kit | 10x Genomics | #PN-1000300 |
| Chromium Next GEM Chip K Automated Single Cell Kit | 10x Genomics | #PN-1000289 |
| Chromium single cell controller | 10x Genomics | #PN-110211 |
| **Software and algorithms** |  |  |
| CLC Genomics Workbench | Qiagen | https://digitalinsights.qiagen.com |
| CellRanger | 10xGenomics | https://support.10xgenomics.com/single-cell-gene-expression/software |
| R | R Foundation for Statistical Computing | https://www.R-project.org |
| Seurat | Satija Lab. | https://github.com/satijalab/seurat |
| SeuratObject | Satija Lab. | https://satijalab.github.io/seurat-object/ |
| Xenium Onboard Analysis | 10x Genomics | SCR_026158 |
| Python programming language | Python Software Foundation | SCR_008394 |
| SciPy | SciPy developers | SCR_008058 |
| Statsmodels | https://www.statsmodels.org | SCR_016074 |
| Matplotlib | Matplotlib developers | SCR_008624 |

### Sample collection, single-cell RNA and TCR-repertoire sequencing

For NSCLC, CRC, BC, and PDAC, sample collection and experimental processing have been described previously (10, 13). In brief, fresh tumor and adjacent normal tissue samples from NSCLC, BC, CRC, and PDAC were selected by a pathologist. These samples were then sliced using scalpels and subjected to combined mechanical and enzymatic tissue dissociation. For NSCLC, CRC, and BC specimens, the Tumor dissociation kit (Miltenyi Biotec, Bergisch-Gladbach, Germany, 130-095-929) in combination with the GentleMACS system (Miltenyi Biotec, 130-093-235) was used according to the manufacturer’s instructions. For BC, this mixture was supplemented with additional *Clostridium histolyticum* collagenase (Sigma Aldrich) at 0.4 mg/ml. For enzymatic dissociation of PDAC, liberase DH (Roche Diagnostics) was used at 25 µg/ml.

Upon filtering through 70 μm-cell strainers (Miltenyi Biotec), aliquots were cryopreserved and the remainder of the cell suspensions was filtered through 30 μm strainers (Miltenyi Biotec), and centrifuged for 7 min at 400 x g to pellet cells after dissociation. NSCLC and CRC samples underwent Percoll gradient centrifugation to enrich for lymphocytes. Cells were resuspended in TexMACS medium (Miltenyi Biotec) supplemented with Penicillin/Streptomycin (Biowest) and allowed to rest overnight. Subsequently, the cells from the overnight cultures were stained using an antibody panel reactive against CD3, CD8, CD4, and PD1. The corresponding isotype control antibodies were used in control stains. CD3^+^ and CD8^+^ leucocyte fractions were isolated from TILs and healthy tissue-infiltrating cells using FACS (BD FACS Melody, Heidelberg, Germany).

Genomic DNAs isolated from sorted TIL fractions and healthy tissue infiltrating lymphocytes as well as from peripheral blood were subjected to TCRSeq as described previously (10). Briefly, genomic DNA from T cell subpopulations was isolated using the QIAamp blood kit (Qiagen, Hilden Germany) and used for the generation of NGS libraries by a two-step PCR protocol (11). In addition, single-cell cDNA-libraries were generated from CD3^+^ or CD8^+^ TIL single-cell suspensions by use of the 10x Genomics® GemCode^TM^ Technology. From this single-cell cDNA, TCR repertoire libraries were generated using the 10x TCR amplification kit and the library construction kit, all v2, according to the manufactureŕs instructions. Following clean-up, TCR repertoire libraries were analyzed by Illumina next generation sequencing in-house on a MiSeq instrument. Single-cell gene expression libraries were analyzed on a HiSeq instrument (StarSEQ GmbH, Mainz, Germany)

### Bioinformatic analysis of single-cell and TCR profiling data

Single-cell gene expression (scGEX) data was generated using the 10x Genomics platform and processed with Cell Ranger v3.1.0 from the previously sorted CD8^+^ T cell fraction (PDAC_49) as well as CD3^+^ T cell fraction (PDAC_10). Downstream analyses, were performed in R v4.3.3 using Seurat v4.4.0 and SeuratObject v4.1.4.

Standard quality-control criteria were applied, requiring a minimum of 500 UMIs and 300 detected genes for PDAC_10 or 250 detected genes for PDAC_49, with a maximum of 20% mitochondrial transcripts per cell. Genes detected in fewer than four cells were excluded. Additionally, all TCR V– and J-segment genes were removed prior to normalization to minimize clustering driven by TCR expression rather than transcriptional cell states.

Single-cell gene expression data were normalized and variance-stabilized using the SCTransform function, with mitochondrial transcript levels regressed out. Principal component analysis (PCA) was performed on the resulting SCT assay, and the first 30 principal components were used for the final UMAP visualization, nearest-neighbor graph construction, and graph-based clustering. Cluster-enriched genes were identified using the FindAllMarkers function with a Wilcoxon rank-sum test, considering only positively enriched genes with a log_2_ fold-change of at least 0.5 and expression in at least 10% of cells. These marker genes were then used for manual cluster annotation based on the guidelines proposed by Masopust et al. (17).

A CD8^+^ T cell re-clustering was performed on the PDAC_10 CD3^+^ scGEX analysis. To do so all cells expressing any CD8 gene, together with cells from previously identified CD8-like clusters (majority of cells with CD8 gene marker) were used. The CD8^+^ subset was re-clustered using the same workflow as described above from the SCTransform function onwards.

Predefined cell populations, including Cluster-A T cells and clonotypes 1–6 from PDAC_10, were mapped onto the corresponding UMAP embeddings using the 10x Genomics cell barcodes to link the single-cell TCR V(D)J sequencing data with the scGEX data.

### Xenium *in situ* analysis

Spatial transcriptomics was performed using the 10x Genomics Xenium Spatial platform with v1 chemistry. FFPE tissue sections were processed according to Document CG000578, including overnight drying, deparaffinization, and decrosslinking as described in the Xenium *In Situ* for FFPE – Deparaffinization & Decrosslinking manual (CG000580). The human multi-tissue and cancer gene expression panel targeting 377 genes was applied together with a 100-target add-on panel and incubated overnight. Post-hybridization washes, ligation, amplification, and cell segmentation labeling were performed following the Xenium Prime *In Situ* Gene Expression manual (CG000760). Xenium slides were processed for imaging and data acquisition following the Xenium Analyzer protocol (CG000584). The instrument operated with instrument software and analysis versions 3.3.0.1.

### Bioinformatic analysis of Xenium spatial transcriptomic data

Cell-level transcript count matrices and spatial coordinates generated by the Xenium platform were analyzed in Python 3.11 using Scanpy and Squidpy together with custom analysis scripts. Custom analysis scripts developed for this study will be made publicly available through GitHub upon publication of the article.

For the detailed transcriptomic and spatial analyses, cell-level quality-control thresholds were selected by visual inspection of transcript-count and detected-gene distributions to exclude cells within the low-count and low-complexity tails. Thresholds were adapted to the respective Xenium dataset. For the direct comparison of PDAC_49 tumor and matched normal pancreas analyzed on the same Xenium slide, identical cell-level quality-control criteria were applied to the complete dataset prior to spatial separation of the two tissue compartments, ensuring directly comparable filtering conditions between tumor and normal tissue. No additional cell-level Q-score filter was applied.

For transcriptome-based cell clustering, transcript counts were normalized to 10,000 counts per cell and log-transformed. Custom specific probes, such as clonotype-specific CDR3 probes, TCR variable-gene probes and mutation-specific probes, were excluded from the feature space used for dimensionality reduction and clustering to prevent cell-state classification from being driven by study-specific tracking features. These probes were retained for dedicated downstream analyses. Principal component analysis (PCA) was followed by neighborhood graph construction using up to 30 principal components, 15 nearest neighbors, and cosine distance, followed by UMAP dimensionality reduction. Cell populations were identified by Leiden community detection at a resolution of 0.5. Cell types and epithelial compartments were annotated based on cluster-enriched marker genes and spatial tissue context.

Cluster-A T cells in PDAC_49, were identified independently of transcriptome-based clustering using custom Xenium probes targeting Cluster-A-associated CDR3α and CDR3β sequences and the characteristic TRBV10-2/TRAV21 gene-segment combination. A Cluster-A TCR signature was assigned when cells showed either paired Cluster-A-associated CDR3α and CDR3β detection, concurrent TRBV10-2 and TRAV21 detection, or a Cluster-A-associated CDR3 signal together with TRBV10-2 or TRAV21. For the detailed PDAC_49 tumor analysis, Cluster-A cells were additionally required to express CD8A or CD8B.

For comparison with Cluster-A CD8⁺ T cells in PDAC_49, a high-confidence non-Cluster-A CD8⁺ T cell population was defined within the transcriptome-defined T cell compartment by expression of CD8A or CD8B together with TCR transcript expression and exclusion of Cluster-A T cells. The T cell compartment was subsequently re-clustered independently, and a naïve/central-memory-like CD8⁺ T cell reference population was defined based on the corresponding transcriptionally annotated subcluster and restricted to high-confidence non-Cluster-A CD8⁺ T cells.

For the direct comparison of PDAC_49 tumor and matched normal pancreas, high-confidence CD8⁺ T cells were defined independently of Leiden clustering by concurrent evidence of leukocyte identity (PTPRC expression), T cell identity (expression of at least one of CD3D, CD3E, CD3G, TRAC, TRBC1-C2, CD247, or CD2), and CD8 expression (CD8A or CD8B). Cluster-A cells were identified using the same CDR3– and TRBV10-2/TRAV21-based criteria described above and were required to fulfill these high-confidence CD8⁺ T cell criteria. The same criteria were applied independently to tumor and matched normal tissue. The proportions of Cluster-A and non-Cluster-A cells among high-confidence CD8⁺ T cells were compared between tumor and matched normal tissue using a two-sided Fisher’s exact test.

For PDAC_10, the six dominant clonotypes (ct1–ct6) identified by single-cell 10x VDJ sequencing were assigned using clonotype-specific CDR3α, CDR3β, TRAV, and TRBV probes. Evidence from these clonotype-specific features was integrated into a feature-based assignment score, with CD8A and CD8B expression included as supportive evidence only when at least one clonotype-specific CDR3 probe was detected in the same cell. Clonotype assignment required a total score of at least three and a uniquely best-supported clonotype. Cells with insufficient or ambiguous evidence remained unassigned. These clonotype assignments were used for the spatial analyses of ct1–ct6.

Spatial proximity was quantified using segmented-cell centroid coordinates. For each clonotype-assigned T cell, the Euclidean distance to the nearest cell belonging to the respective annotated epithelial compartment was calculated using nearest-neighbor searches. In PDAC_49, distances were determined to the individual malignant epithelial clusters, their combined malignant compartment, and the non-malignant ductal epithelial compartment. Distance distributions of Cluster-A CD8⁺ T cells and naïve/central-memory-like CD8⁺ T cells were compared using two-sided Mann–Whitney U tests, with Benjamini–Hochberg correction for multiple comparisons. To assess whether the unequal numbers of T cells in these two populations influenced statistical significance, naïve/central-memory-like CD8⁺ T cells were randomly subsampled to match the number of Cluster-A CD8⁺ T cells with valid distance measurements across all three malignant compartments. This matched-size analysis was repeated 1,000 times, with Mann–Whitney U testing and Benjamini–Hochberg correction performed independently for each iteration. Distances of Cluster-A CD8⁺ T cells to malignant and non-malignant ductal epithelial compartments were compared descriptively because these epithelial compartments differed in abundance and spatial distribution.

In PDAC_10, nearest-cell distances of ct1–ct6 were determined for the invasive adenocarcinoma compartment, the intraepithelial neoplasia compartment, and their combined neoplastic epithelial compartment. Tumor-to-normal (T/N) ratios for ct1–ct6 were determined independently by bulk TCRβ repertoire sequencing of CD8⁺ T cells isolated from matched tumor and adjacent normal tissue and were calculated from the relative clonotype frequencies in the respective TCR repertoires. Clonotypes were ordered according to decreasing T/N ratio for comparison of their spatial distributions and transcriptional features. Associations between T/N ratios and clonotype-level marker-positive cell fractions were assessed using two-sided Spearman rank correlations. Where multiple hypothesis tests were performed within an analysis, P values were adjusted using the Benjamini–Hochberg false-discovery rate procedure. Unless otherwise stated, statistical tests were two-sided.

### Epstein–Barr virus-encoded RNA *in situ* hybridization

To detect Epstein–Barr virus (EBV)-encoded RNA (EBER), we performed an *in situ* hybridization (ISH) assay as previously described (18). Formalin-fixed, paraffin embedded tissue blocks were cut into 3 µm thin sections. A ready-to-use EBER-ISH probe (BOND #PB5089) was used together with the Leica Bond-maX autostainer (Leica Biosystems, Illinois, USA) according to the standard BOND protocol.

## Results

Complementing our established workflow, we explored whether Xenium spatial transcriptomics could provide orthogonal *in situ* evidence supporting the tumor association of tumor-enriched TCR clonotypes by linking clonotype identity to cellular state and tissue architecture at single-cell resolution. We therefore integrated tumor enrichment, spatial localization, and transcriptional phenotype to characterize selected candidate tumor-reactive TCR clonotypes (Fig. 1).

### Cluster-A TCRs are enriched in tumor tissue and detectable in peripheral blood

Cluster-A TCRs represent a previously identified and characterized group of public sequences detected in a subgroup of HLA-A*02-positive patients with different types of solid cancers (13). Several lines of evidence connect some of these TCRs with reactivity against EBV (19–22). However, TCR-T cells genetically engineered to express representative examples of these TCRs recognized a set of HLA-A*02:01-positive tumor cell lines with no evidence of EBV-infection, suggesting cross reactivity with a shared tumor-associated antigen (13, 16, 21). To systematically assess the distribution of Cluster-A T cells across tumor, adjacent non-tumor tissue, and peripheral blood, we analyzed Cluster-A TCR frequencies in paired samples across 14 solid tumor cases, comprising six non-small cell lung cancer (NSCLC), four pancreatic ductal adenocarcinoma (PDAC), two breast cancer (BC), and two colorectal cancer (CRC) cases (Table 2). Consistent with our previous report, Cluster-A TCRs generally showed preferential enrichment in tumor tissue compared with matched adjacent non-tumor tissue (13). Cluster-A TCRs were also consistently detectable at low frequencies in matched peripheral blood samples across tumor entities (Table 2). Independent reanalysis of previously published NSCLC TCRβ repertoire data further supported this tissue distribution pattern (Table S1).

**Table 2:**
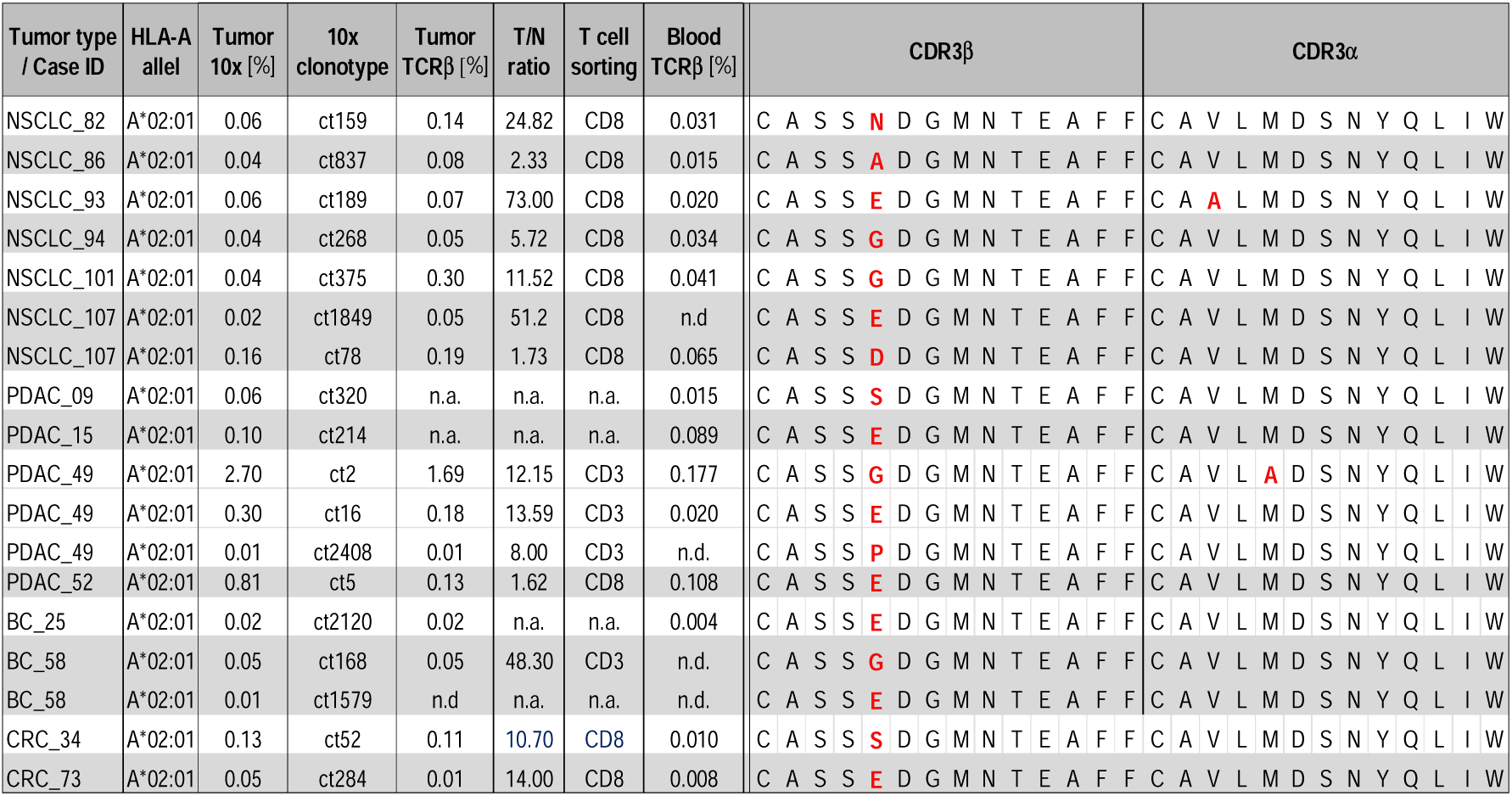
Distribution, frequencies, and sequence characteristics of Cluster-A TCR clonotypes across tumor entities. Cluster-A TCRs were defined by CDR3 sequence homology and usage of TRBV10-2, TRAV21, TRBJ1-1, and TRAJ33. All 14 patients harboring Cluster-A TCRs carried the HLA-A*02:01 allele. Tumor/non-tumor (T/N) ratios were determined by TCR profiling of CD3^+^– or CD8^+^-sorted T cells (see “T cell sorting” column). Tumor 10x [%] indicates the frequency of each clonotype among T cells identified by 10x single-cell V(D)J sequencing, whereas Tumor TCRβ [%] and Blood TCRβ [%] indicate clonotype frequencies determined by bulk TCRβ repertoire sequencing in tumor and peripheral blood, respectively. A subset of cases (NSCLC_82, NSCLC_86, NSCLC_93, NSCLC_94, NSCLC_101, and BC_25) overlaps with previously published datasets reporting tumor enrichment of Cluster-A TCRs; peripheral blood was not assessed in the previous study (13). For T/N ratios where the non-tumor value was zero, the non-tumor frequency was uniformly set to 0.001%. n.a., not analyzed; n.d., not detected; ct, clonotype.

| Tumor type / Case ID | HLA-A allel | Tumor 10x [%] | 10x clonotype | Tumor TCRβ [%] | T/N ratio | T cell sorting | Blood TCRβ [%] | CDR3β | CDR3α |
|---|---|---|---|---|---|---|---|---|---|
| NSCLC_82 | A*02:01 | 0.06 | ct159 | 0.14 | 24.82 | CD8 | 0.031 | CASS**N**DGMNTEAFF | CAVLMDSNYQLIW |
| NSCLC_86 | A*02:01 | 0.04 | ct837 | 0.08 | 2.33 | CD8 | 0.015 | CASS**A**DGMNTEAFF | CAVLMDSNYQLIW |
| NSCLC_93 | A*02:01 | 0.06 | ct189 | 0.07 | 73.00 | CD8 | 0.020 | CASS**E**DGMNTEAFF | CA**A**LMDSNYQLIW |
| NSCLC_94 | A*02:01 | 0.04 | ct268 | 0.05 | 5.72 | CD8 | 0.034 | CASS**G**DGMNTEAFF | CAVLMDSNYQLIW |
| NSCLC_101 | A*02:01 | 0.04 | ct375 | 0.30 | 11.52 | CD8 | 0.041 | CASS**G**DGMNTEAFF | CAVLMDSNYQLIW |
| NSCLC_107 | A*02:01 | 0.02 | ct1849 | 0.05 | 51.2 | CD8 | n.d | CASS**E**DGMNTEAFF | CAVLMDSNYQLIW |
| NSCLC_107 | A*02:01 | 0.16 | ct78 | 0.19 | 1.73 | CD8 | 0.065 | CASS**D**DGMNTEAFF | CAVLMDSNYQLIW |
| PDAC_09 | A*02:01 | 0.06 | ct320 | n.a. | n.a. | n.a. | 0.015 | CASS**S**DGMNTEAFF | CAVLMDSNYQLIW |
| PDAC_15 | A*02:01 | 0.10 | ct214 | n.a. | n.a. | n.a. | 0.089 | CASS**E**DGMNTEAFF | CAVLMDSNYQLIW |
| PDAC_49 | A*02:01 | 2.70 | ct2 | 1.69 | 12.15 | CD3 | 0.177 | CASS**G**DGMNTEAFF | CAVL**A**DSNYQLIW |
| PDAC_49 | A*02:01 | 0.30 | ct16 | 0.18 | 13.59 | CD3 | 0.020 | CASS**E**DGMNTEAFF | CAVLMDSNYQLIW |
| PDAC_49 | A*02:01 | 0.01 | ct2408 | 0.01 | 8.00 | CD3 | n.d. | CASS**P**DGMNTEAFF | CAVLMDSNYQLIW |
| PDAC_52 | A*02:01 | 0.81 | ct5 | 0.13 | 1.62 | CD8 | 0.108 | CASS**E**DGMNTEAFF | CAVLMDSNYQLIW |
| BC_25 | A*02:01 | 0.02 | ct2120 | 0.02 | n.a. | n.a. | 0.004 | CASS**E**DGMNTEAFF | CAVLMDSNYQLIW |
| BC_58 | A*02:01 | 0.05 | ct168 | 0.05 | 48.30 | CD3 | n.d. | CASS**G**DGMNTEAFF | CAVLMDSNYQLIW |
| BC_58 | A*02:01 | 0.01 | ct1579 | n.d | n.a. | n.a. | n.d. | CASS**E**DGMNTEAFF | CAVLMDSNYQLIW |
| CRC_34 | A*02:01 | 0.13 | ct52 | 0.11 | 10.70 | CD8 | 0.010 | CASS**S**DGMNTEAFF | CAVLMDSNYQLIW |
| CRC_73 | A*02:01 | 0.05 | ct284 | 0.01 | 14.00 | CD8 | 0.008 | CASS**E**DGMNTEAFF | CAVLMDSNYQLIW |

### In-depth analysis of Cluster-A TCRs in PDAC

For integrated single-cell and spatial transcriptomic analyses, we selected PDAC_49, in which three distinct Cluster-A TCRs were detected among CD3⁺ tumor-infiltrating lymphocytes (TILs). We first characterized these clonotypes individually before analyzing Cluster-A T cells collectively at the transcriptional and spatial levels.

In PDAC_49 TILs, we identified 242 T cells with paired Cluster-A TCRα/β clonotypes, corresponding to the three distinct CDR3α/β amino acid sequence pairs. Table 2 shows the CDR3 sequences of the clonotypes and summarizes the T/N ratio in bulk CD3⁺ T cells. Beyond the tabulated data, 10x single-cell V(D)J sequencing of CD8⁺ TILs revealed that Cluster-A clonotype PDAC_49_ct2 was the second most abundant clonotype, with a relative clonotype frequency of 2.7%. This dominant clonotype showed a positive T/N ratio of 12× in bulk CD3⁺ T cells and was further enriched 19× in bulk CD3⁺/PD1⁺ compared to CD3⁺/PD1⁻ T cells, representing 1% of the entire PD1⁺ CD3⁺ compartment. This phenotype is characteristic of activated/chronically stimulated TILs.

### Single-cell analysis identifies Cluster-A clonotypes as activated cytotoxic CD8⁺ effector T cells

Single-cell gene expression profiling revealed Cluster-A T cells as a transcriptionally distinct population of activated cytotoxic CD8⁺ T cells, predominantly mapped to tissue effector memory (T-EM) accompanied by smaller fractions of tissue-resident memory (T-RM) cells and few central memory (T-CM) cells. This distribution is consistent with a differentiation continuum across T-CM, T-RM, and T-EM states, potentially reflecting different stages of antigen experience, activation, and effector differentiation within the tumor (Fig. 2). In PDAC_49, Cluster-A T cells predominantly mapped to the T-EM compartment and displayed an activated effector phenotype characterized by expression of cytotoxic and effector-associated markers, including GZMA, GZMK, KLRG1, accompanied by CST7 expression (Fig. 2).

**Figure 2:**
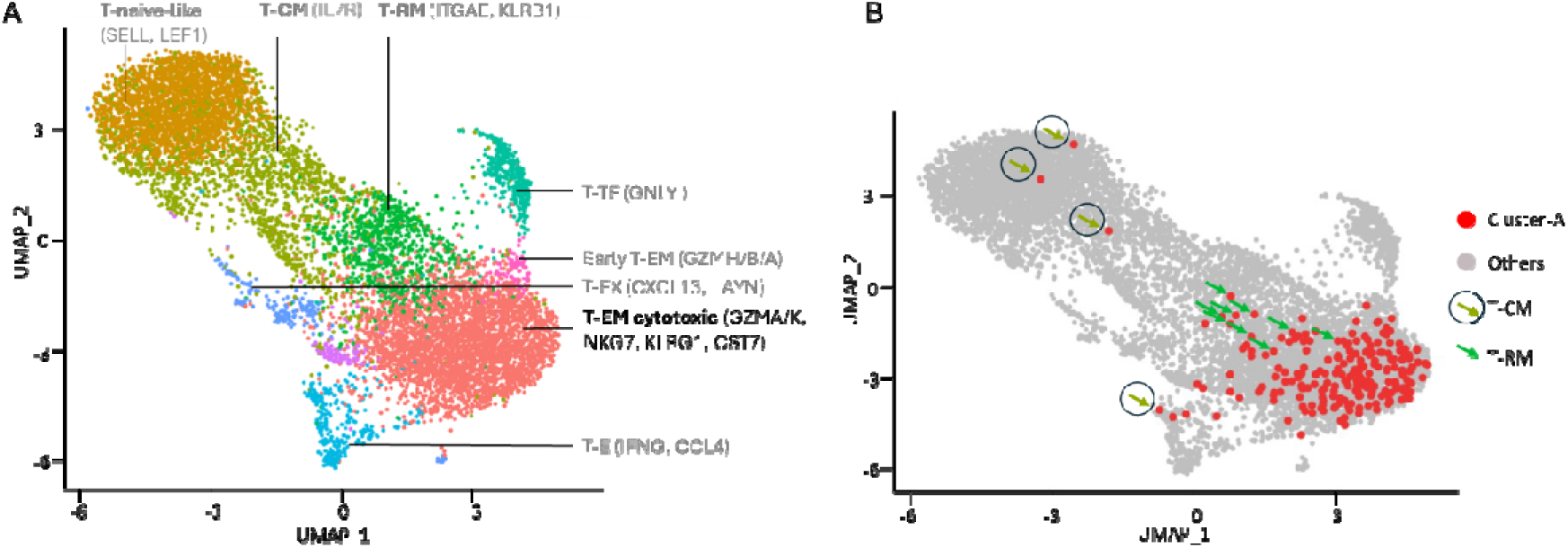
UMAP analysis of CD8⁺ Cluster-A T cell clonotypes. (**A**) UMAP projection of CD8⁺ T cells from PDAC_49. (**B**) Cluster-A CD8⁺ T cells localized via cellular barcodes and highlighted in red. Cluster-A clonotypes are predominantly composed of effector memory T cells (T-EM), with smaller fractions of tissue-resident memory T cells (T-RM; green arrows) and central memory T cells (T-CM; olive-green arrows), consistent with a differentiation continuum across antigen-experienced CD8⁺ T cell states. These distinct antigen-experienced T cell states may reflect different stages of activation and effector differentiation. T cell subset nomenclature follows the guidelines proposed by David Masopust et al. (17).

### Xenium spatial analysis demonstrates tumor-restricted localization of Cluster-A CD8 T cells in PDAC_49

To spatially validate the tumor-associated enrichment of Cluster-A T cells in whole tissue context, we compared their distribution between PDAC_49 tumor tissue and matched adjacent normal pancreas using Xenium spatial transcriptomics. Cluster-A T cells were identified using a stringent Xenium-compatible signature requiring Cluster-A-specific TCR features (Cluster-A-associated CDR3 transcripts and the characteristic TRBV10-2/TRAV21 gene segment combination) together with CD8 marker expression. High-confidence CD8⁺ T cells were independently defined using a hierarchical marker strategy requiring lymphocyte, T cell, and CD8 marker expression. All high-confidence CD8⁺ T cells not classified as Cluster-A were used as the reference population to compare the relative frequency of Cluster-A cells between tumor and matched normal tissue.

The Xenium experiment contained tumor tissue and matched normal pancreas on the same slide, enabling direct spatial comparison under identical experimental conditions. We identified a total of 13,219 CD8⁺ T cells in tumor tissue, comprising 62 Cluster-A and 13,157 other high-confidence CD8⁺ T cells, compared with 1,720 high-confidence CD8⁺ T cells in matched normal pancreas, none of which were classified as Cluster-A. Applying the same Cluster-A classification strategy to both compartments demonstrated significant tumor-associated enrichment of Cluster-A CD8⁺ T cells (62 vs. 13,157 in tumor; 0 vs. 1,720 in normal tissue; two-sided Fisher’s exact test, *P* < 0.001).

The absence of Cluster-A cells in normal pancreas was not attributable to insufficient CD8⁺ T cell representation, as 1,720 high-confidence CD8⁺ T cells were detected in the matched normal tissue. Spatial visualization confirmed that Cluster-A cells were exclusively detected within the tumor compartment, whereas high-confidence CD8⁺ T cells lacking the Cluster-A signature were present in both tumor and normal pancreas (Fig. 3).

**Figure 3:**
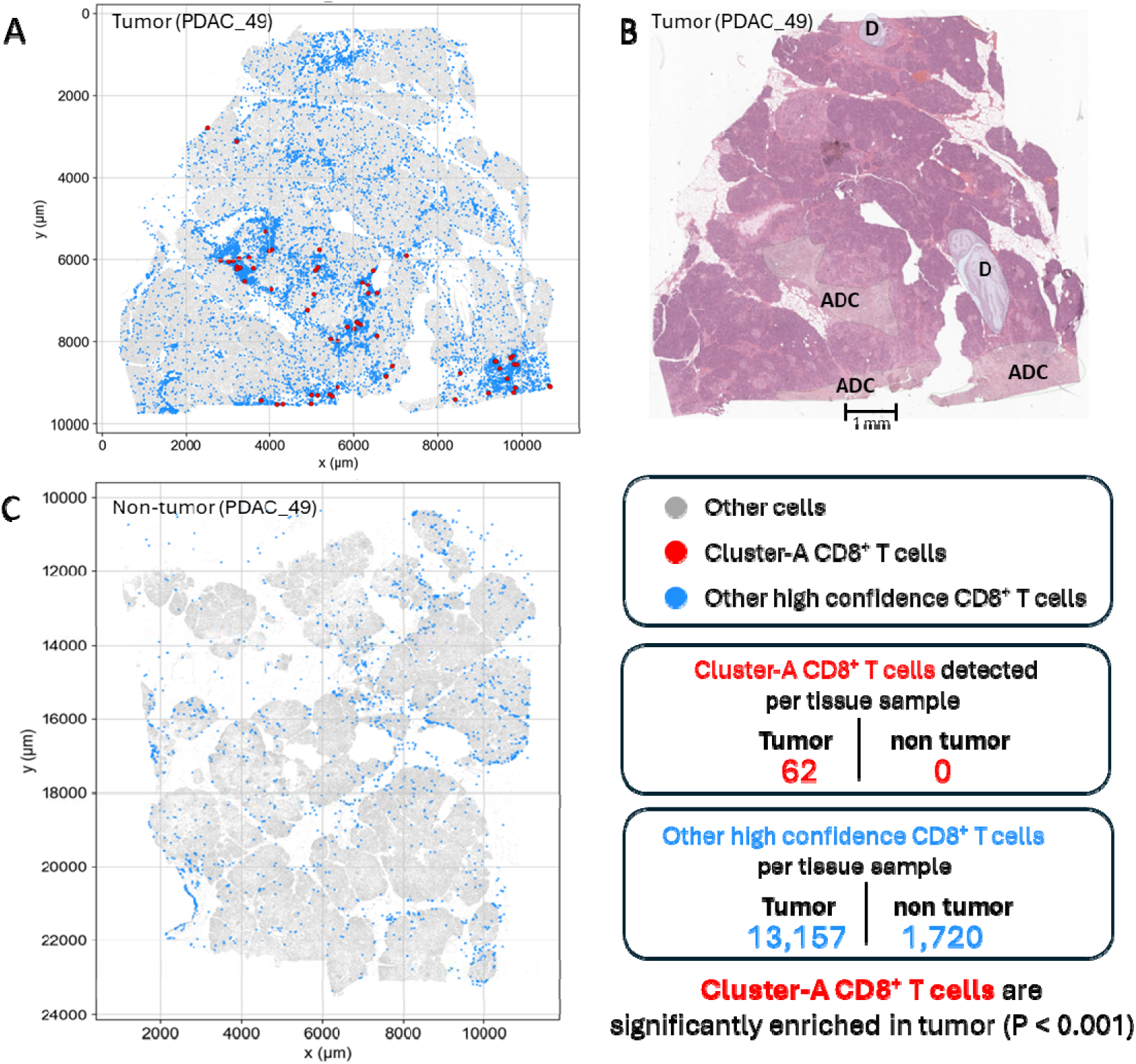
Xenium spatial analysis demonstrates tumor-restricted localization of Cluster-A CD8^+^ T cells in PDAC_49. (**A**) Spatial distribution of high-confidence CD8^+^ T cells (blue) and Cluster-A CD8 T cells (red) in the tumor region of the PDAC_49 Xenium section. All segmented cells are shown in light gray. Cluster-A CD8 T cells localize within the adenocarcinoma compartment annotated according to Hematoxylin and eosin (H&E) staining. (**B**) H&E staining of the same Xenium slide, performed after completion of the Xenium spatial transcriptomic analysis. Adenocarcinoma regions (ADC) and pancreatic ducts (D) were annotated by a pathologist. (**C**) Spatial distribution of high-confidence CD8^+^ T cells in the matched normal pancreas. Cluster-A CD8⁺ T cells were detected exclusively in tumor tissue (62 Cluster-A versus 13,157 other high-confidence CD8⁺ T cells) and were absent from matched normal pancreas (0 Cluster-A versus 1,720 other high-confidence CD8⁺ T cells), demonstrating significant tumor-associated enrichment (two-sided Fisher’s exact test, *P* < 0.001).

This finding from spatial analysis was independently supported by TCRβ repertoire analysis of TILs and normal tissue-infiltrating lymphocytes of the same patient, which likewise failed to detect Cluster-A clonotypes in matched normal pancreas. Together, these orthogonal analyses demonstrate that Cluster-A CD8^+^ T cells were restricted to the tumor compartment and were undetectable in the corresponding non-tumor tissue.

### Cluster-A CD8^+^ T cells preferentially localize near KRAS_G12V-positive malignant epithelial compartments and within intratumoral TLS

Having established the tumor-restricted localization of Cluster-A CD8⁺ T cells by direct comparison with matched normal pancreas, we next asked whether Cluster-A cells also exhibit preferential spatial positioning relative to malignant epithelial cells within the tumor microenvironment. To molecularly define malignant epithelial compartments beyond the histopathological tumor annotation used for the tumor-versus-normal comparison, we analyzed an independent PDAC_49 Xenium dataset, integrating histopathological annotation with unsupervised cell-state classification and mutation-specific genetic validation.

Histopathological examination of the corresponding Hematoxylin and Eosin (H&E) stained section identified moderately differentiated (ADC G2) and poorly differentiated (ADC G3) adenocarcinoma regions, together with pancreatic ducts and intratumoral tertiary lymphoid structures (TLS) (Fig. 4A). Unsupervised Leiden clustering of the Xenium transcriptomic data identified two epithelial clusters (clusters 3 and 4) with transcriptional programs characteristic of pancreatic ductal adenocarcinoma, whereas distinct epithelial populations represented non-malignant ductal epithelium (cluster 9) and endocrine pancreatic cells (cluster 10) (Fig. 4B, C). The mutation-specific KRAS_G12V probe was intentionally excluded from the clustering procedure and mapped only after cluster annotation, thereby providing independent genetic validation of malignant epithelial identity. KRAS_G12V transcripts localized to malignant epithelial clusters 3 and 4, supporting their classification as the major malignant epithelial compartments within the tissue (Fig. 4C).

**Figure 4:**
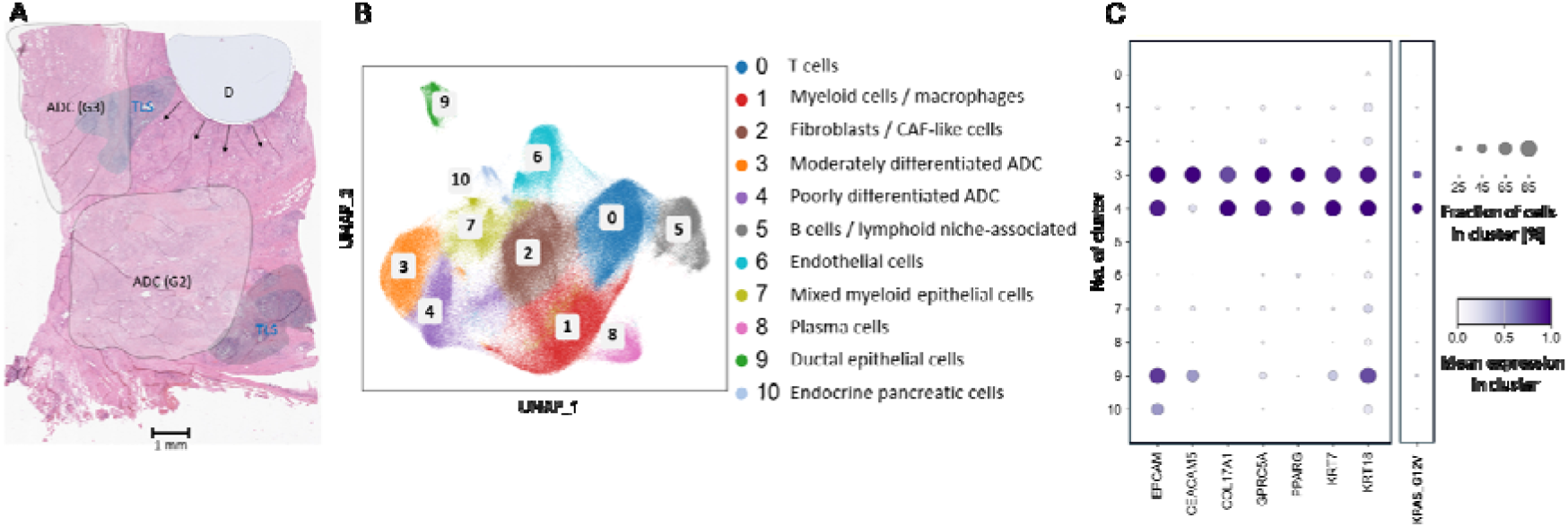
Histopathological and transcriptomic identification of genetically validated malignant epithelial compartments in PDAC_49. (**A**) H&E staining of the PDAC_49 tissue section following Xenium spatial transcriptomic analysis. A large pancreatic duct (D) and radiating smaller ducts with intraepithelial neoplasia (IEN; arrows) are indicated. Regions of moderately differentiated adenocarcinoma (ADC, grade 2 (G2)), poorly differentiated adenocarcinoma (ADC, grade 3 (G3)), and tertiary lymphoid structures (TLS) are annotated. Scale bar, 1 mm. (**B**) UMAP visualization of cell states identified by unsupervised Leiden clustering of the PDAC_49 Xenium dataset. Two epithelial populations were annotated as malignant epithelial clusters (clusters 3 and 4), whereas cluster 9 represented non-malignant ductal epithelium and cluster 10 endocrine pancreatic cells. The remaining clusters correspond to T cells, fibroblasts/CAF-like cells, myeloid cells, mixed epithelial–myeloid cells, endothelial cells, B-cell/lymphoid aggregates and plasma cells. (**C**) Dot plot showing representative markergene expression across Leiden clusters. Malignant epithelial clusters 3 and 4 display complementary epithelial transcriptional programs, whereas cluster 9 represents non-malignant ductal epithelium. The mutation-specific KRAS_G12V probe was intentionally excluded from the clustering procedure and mapped only after cluster annotation, providing independent genetic validation of the malignant epithelial compartments.

Following molecular and genetic definition of the malignant epithelial compartments, we next examined the spatial relationships of Cluster-A CD8 T cells relative to cells within these tumor regions. Compared with an independently defined naïve/central-memory-like CD8 T cell reference population derived from unsupervised subclustering of the global T cell compartment, Cluster-A CD8 T cells localized markedly closer to malignant cluster 3 (190 versus 499 μm) and malignant cluster 4 (191 versus 523 μm) (both FDR-adjusted *P* < 0.0001; Fig. 5A), as well as to the combined malignant compartment (median distance, 168 versus 477 μm; FDR-adjusted *P* < 0.0001). As a sensitivity analysis, the naïve/central-memory-like CD8⁺ T cell reference population was repeatedly subsampled to match the number of Cluster-A CD8⁺ T cells (n = 555 per group). Across 1,000 independent resampling iterations, Cluster-A CD8⁺ T cells remained significantly closer to all malignant epithelial compartments in 100% of iterations (all FDR < 0.0001).

**Figure 5:**
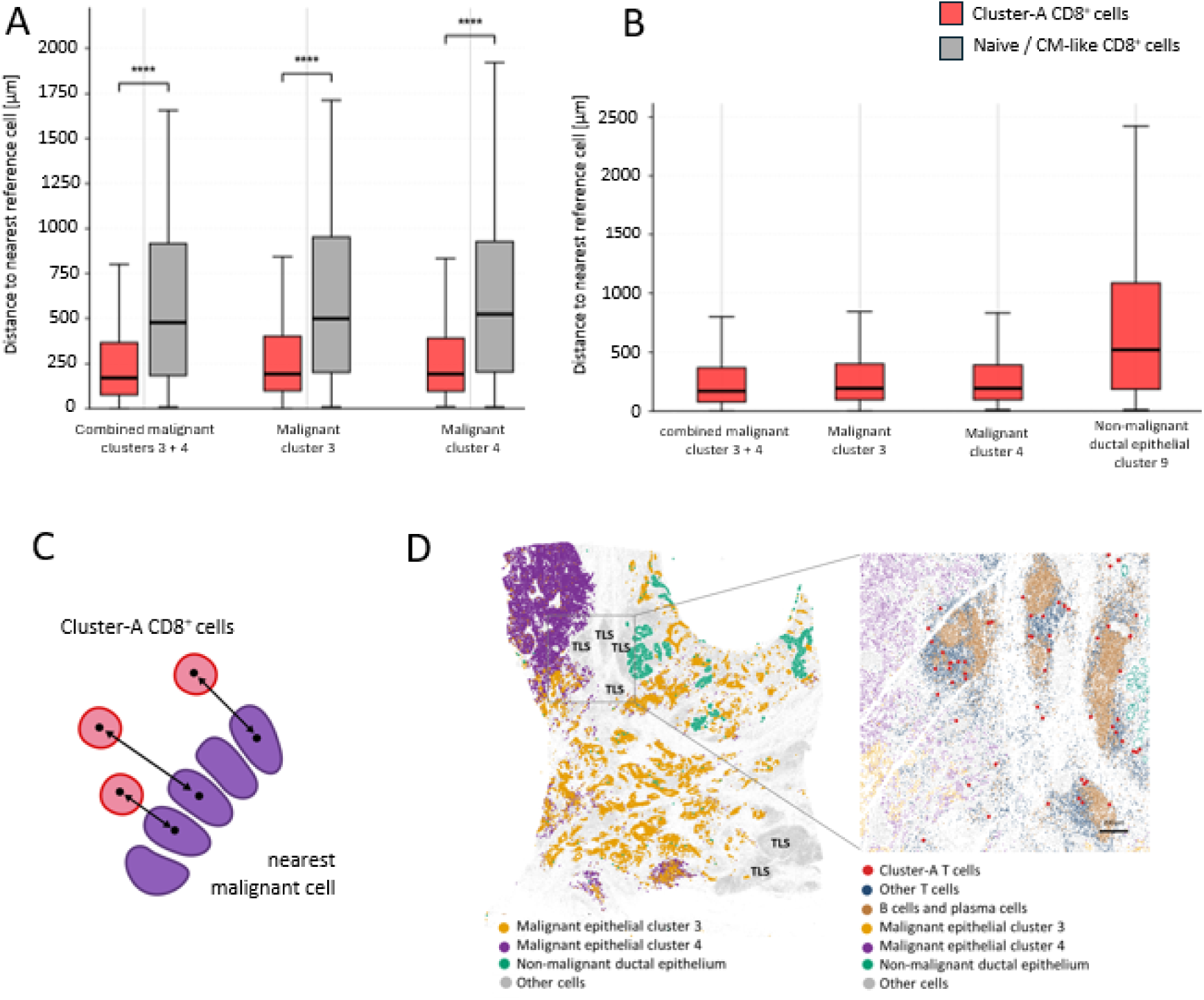
Spatial localization and quantitative evaluation of tumor proximity of Cluster-A CD8⁺ T cells in PDAC_49. (**A**) Nearest-cell distances of Cluster-A CD8⁺ T cells (n = 555) and T cell subcluster-defined naive/central memory (CM)-like CD8^+^ T cell (n = 4,915) to cells of the combined malignant epithelial compartment (clusters 3+4) and to clusters 3 and 4 individually. Cluster-A CD8⁺ T cells localized significantly closer to all malignant compartments (median distance: 168 vs 477 µm for clusters 3+4; 190 vs 499 µm for cluster 3; 191 vs 523 µm for cluster 4; all FDR-adjusted *P* < 0.0001). (**B**) Distances of Cluster-A CD8⁺ T cells to malignant epithelial compartments and non-malignant ductal epithelial cluster 9. Cluster-A cells showed shorter median distances to the combined malignant compartment (clusters 3+4; 168 µm), cluster 3 (190 µm), and cluster 4 (191 µm) than to non-malignant ductal epithelium (518 µm). This comparison is descriptive, as the malignant and non-malignant epithelial compartments differ in abundance and spatial distribution. Boxes in A and B indicate interquartile ranges with median lines; whiskers extend to 1.5 × IQR. Comparisons in A used two-sided Mann– Whitney U tests. (**C**) Schematic illustration of the nearest-cell distance calculation used in A and B. For each Cluster-A CD8⁺ T cell (red), the Euclidean distance between cell centroids was calculated to the nearest cell of the respective epithelial compartment (purple). (**D**) Spatial projection of annotated epithelial and immune cell populations onto the PDAC_49 Xenium section. Malignant epithelial clusters 3 and 4 represent moderately differentiated (G2) and poorly differentiated (G3) tumor compartments, respectively, whereas non-malignant ductal epithelial cluster 9 occupies anatomically distinct regions. The magnified region shows intra– and peritumoral tertiary lymphoid structures (TLS) adjacent to malignant epithelium, with Cluster-A CD8⁺ T cells (red) detected among other T cells (dark blue) and B cells/plasma cells (brown). Other cells are shown in grey. Scale bars, 200 µm.

Consistent with this tumor-proximal localization, Cluster-A CD8⁺ T cells showed shorter median distances to the malignant epithelial compartments (168–191 μm) than to the non-malignant ductal epithelial compartment (518 μm), although this comparison was considered descriptive because the compartments differed in abundance and spatial distribution (Fig. 5B).

To further explore the spatial context of Cluster-A CD8⁺ T cells, we examined their presence within tertiary lymphoid structures (TLS). High-resolution analysis of histopathologically identified TLS revealed multiple Cluster-A CD8⁺ T cells embedded among T cells, B cells, and plasma cells. TLS were located both intratumorally and peritumorally (Fig. 5D).

Together, these analyses demonstrate preferential spatial positioning of Cluster-A CD8 T cells toward genetically validated malignant epithelial compartments within PDAC_49.

To test whether the previously described association of Cluster-A clonotypes with EBV could explain their accumulation in tumor tissue, EBER *in situ* hybridization was performed. No EBV-positive cells were detected in PDAC_49, arguing against ongoing local EBV infection as the driver of Cluster-A accumulation (Supplementary Fig. S1).

### Cluster-A CD8^+^ T cells exhibit a conserved effector/T-EM-associated transcriptional program

To further characterize the transcriptional state of Cluster-A CD8^+^ T cells, we compared clonotype-defined Cluster-A cells with naïve/central-memory-like CD8^+^ T cell reference populations in independent scRNA-seq and Xenium analyses of PDAC_49. Across both modalities, Cluster-A cells exhibited a concordant effector/T-EM-associated transcriptional profile, with increased representation of *GZMA*, *GZMK*, and *CCL5*, whereas *TCF7*, *CCR7*, and *SELL* were preferentially expressed in the respective naïve/central-memory-like populations (Fig. 6). *NKG7*, *PRF1*, *CXCR3*, *KLRG1*, and *CD27* were detected in subsets of Cluster-A cells, consistent with transcriptional heterogeneity within an effector-memory-associated state rather than a uniformly terminally differentiated phenotype. Importantly, in both analyses, the naïve/central-memory-like reference populations were identified by unsupervised clustering of the T cell compartment rather than by selection for the markers subsequently examined, while Cluster-A identity was defined independently by TCR clonotype. Thus, independent single-cell and spatial transcriptomic analyses consistently support a predominantly effector/T-EM-associated phenotype of Cluster-A CD8^+^ T cells.

**Figure 6:**
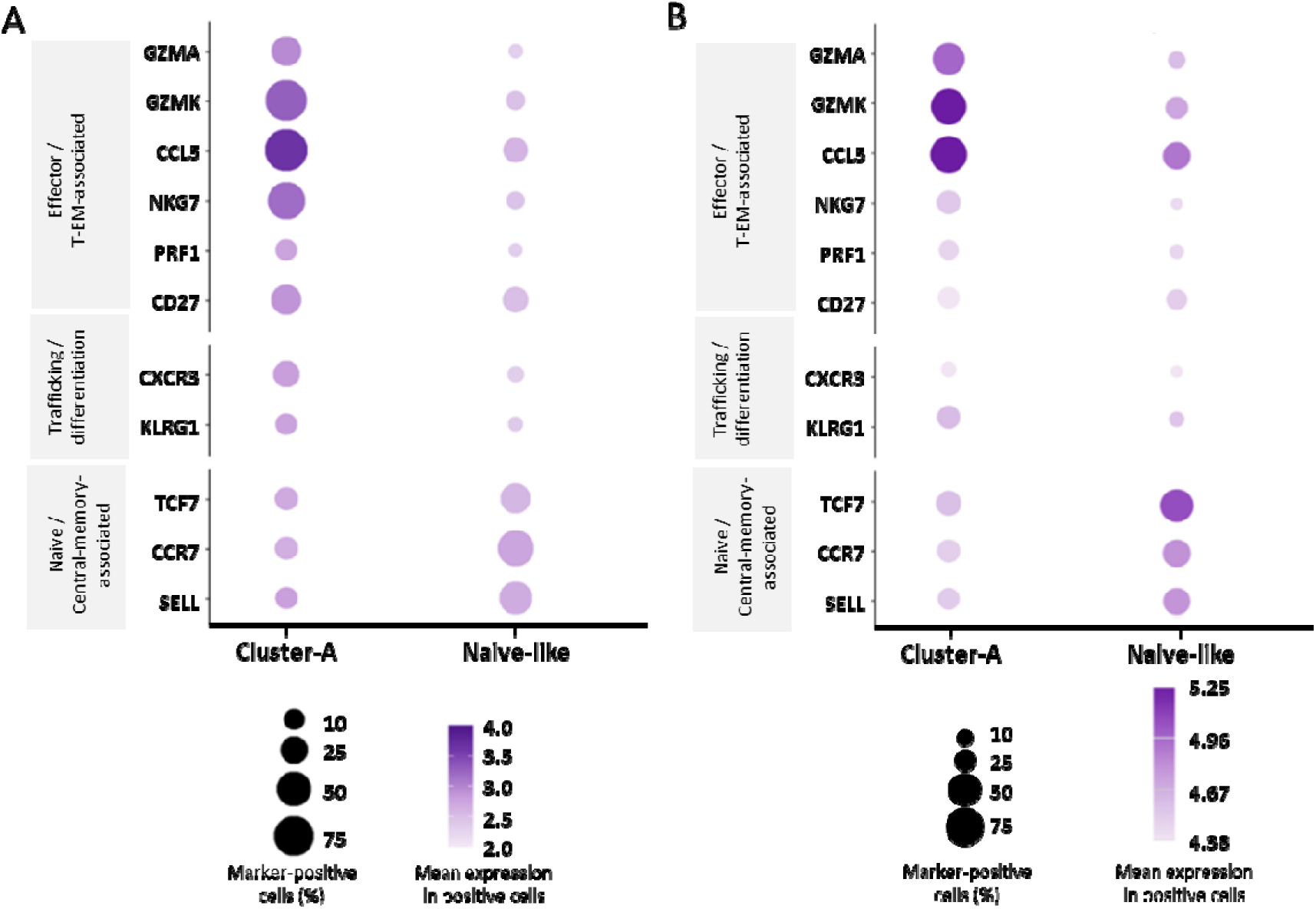
Conserved effector/T-EM-associated transcriptional program of Cluster-A CD8 T cells across single-cell and spatial transcriptomic analyses. (**A**) Single-cell RNA sequencing (scRNA-seq) analysis of PDAC_49 comparing clonotype-defined Cluster-A CD8^+^ T cells with a naïve/central-memory-like CD8^+^ T cell population identified by unsupervised clustering of the T cell compartment. (**B**) Xenium spatial transcriptomic analysis of PDAC_49 comparing clonotype-defined Cluster-A CD8 T cells with naïve/central-memory-like CD8 T cells identified by unsupervised T cell subclustering and restricted to high-confidence non-Cluster-A CD8 T cells. Cluster-A T cell identity was defined independently of transcriptional clustering, based on their characteristic TCR sequence features. Markers are grouped into effector/T-EM-(GZMA, GZMK, CCL5, NKG7, PRF1, CD27), trafficking/differentiation-(CXCR3, KLRG1), and naïve/central-memory-associated (TCF7, CCR7, SELL) genes. Dot size indicates the percentage of marker-positive cells; color indicates mean normalized expression among marker-positive cells. Expression scales are modality-specific.

### Tumor enrichment is associated with cytotoxic effector-memory states across independent clonotypes in PDAC_10

After establishing that the recurrent Cluster-A T cell population is enriched in tumor tissue, exhibits an effector-associated phenotype, and preferentially localizes in proximity to malignant epithelial cells, we next asked whether these features extend beyond this shared clonotype family. To address this question in an independent setting, we analyzed the six most dominant CD8⁺ T cell clonotypes (ct1–ct6) from PDAC_10 for which tumor enrichment, transcriptional state, and spatial localization could be assessed independently. Tumor-to-normal (T/N) ratios had previously been determined by bulk TCRβ repertoire sequencing of CD8⁺ T cells isolated from matched tumor and adjacent normal tissue, providing a compartment-level measure of clonotype enrichment independent of single-cell transcriptional and spatial information. The clonotypes spanned a broad range of tumor enrichment, from strongly tumor-enriched ct5 (T/N = 21.25), ct2 (T/N = 7.17), and ct4 (T/N = 4.05) to non-enriched ct1 (T/N = 0.82), ct3 (T/N = 0.34), and ct6 (T/N = 0.05).

We first examined whether this differential tumor enrichment was associated with distinct CD8⁺ T cell transcriptional states. Single-cell RNA and paired TCR sequencing resolved major CD8⁺ T cell differentiation states, including naïve-like/central memory (T-CM), tissue-resident memory (T-RM), cytotoxic effector-memory (T-EM), effector-memory (T-EM) and terminal effector (T-TE) states (Fig. 7A). Projection of ct1–ct6 onto this transcriptional landscape revealed clonotype-specific distributions (Fig. 7B). The tumor-enriched clonotypes ct5, ct2, and ct4 preferentially localized to cytotoxic T-EM and T-EM states, whereas clonotypes with low T/N ratios showed broader and distinct distributions across the transcriptional landscape.

**Figure 7:**
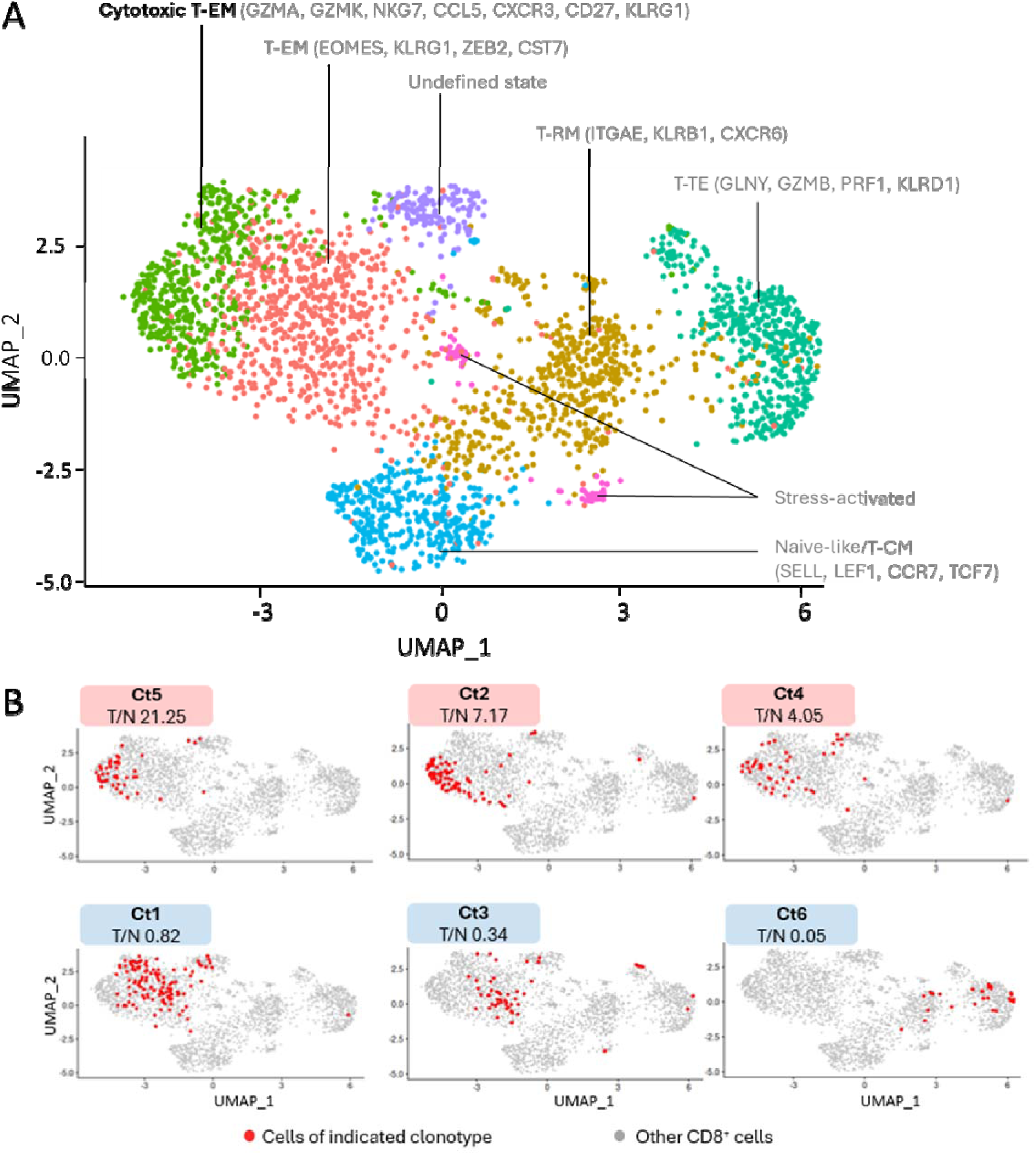
Single-cell profiling links tumor enrichment to effector differentiation in PDAC_10. (**A**) UMAP of CD8⁺ T cells annotated according to major transcriptional states, including naïve-like/central memory (T-CM), tissue-resident memory (T-RM), effector-memory (T-EM), cytotoxic T-EM, terminal effector (T-TE), stress-activated, and undefined states. Selected marker genes used to characterize the respective transcriptional states are indicated. (**B**) UMAP projections of the six most dominant PDAC_10 CD8⁺ T cell clonotypes (ct1–ct6), with clonotype-assigned cells highlighted in red against the overall CD8⁺ T cell landscape in grey. Clonotypes are ordered by decreasing tumor-to-normal (T/N) ratio, as determined by bulk TCRβ repertoire sequencing of matched tumor and adjacent normal tissue. Clonotypes with T/N ratios > 4 are indicated by light-red labels, whereas clonotypes with T/N ratios < 1 are indicated by light-blue labels.

### Tumor enrichment tracks spatial proximity to neoplastic epithelial compartments in PDAC_10

We next asked whether this association between tumor enrichment and cytotoxic effector-memory differentiation was also reflected in the spatial organization of these clonotypes within the tumor tissue. Xenium spatial transcriptomics of PDAC_10 identified distinct epithelial compartments corresponding to invasive adenocarcinoma (cluster 3), intraepithelial neoplasia (cluster 8), and non-malignant pancreatic epithelium (clusters 5 + 6) (Fig. 8A–D). We then mapped ct1–ct6 onto the tissue section and quantified their distances to invasive adenocarcinoma (cluster 3), intraepithelial neoplasia (cluster 8), and the combined neoplastic epithelial compartment (clusters 3 + 8).

**Figure 8:**
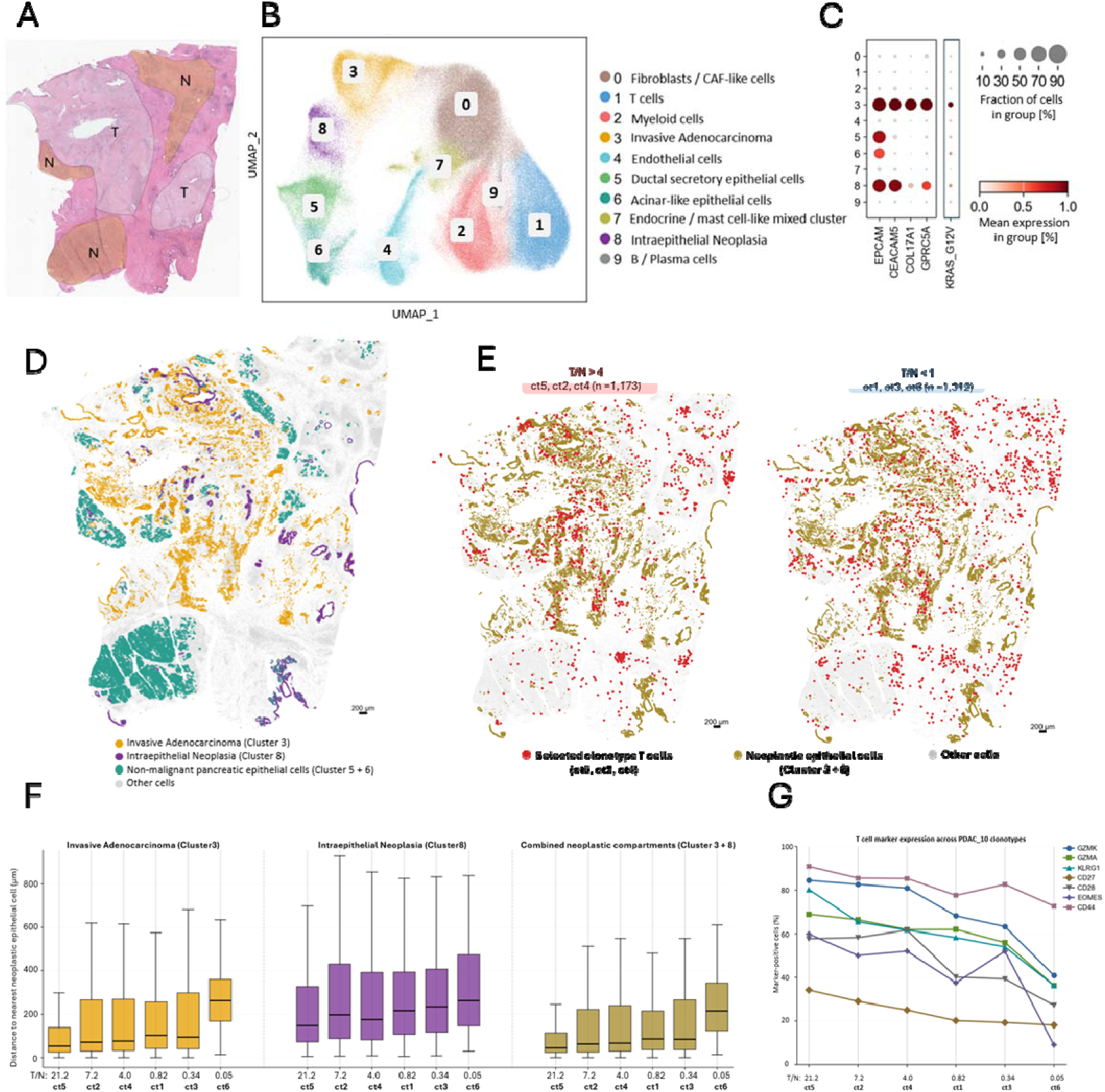
Tumor enrichment across independent CD8⁺ T cell clonotypes is associated with spatial tumor proximity and effector differentiation in PDAC_10. (**A**) Representative H&E-stained section of the PDAC_10 tissue analyzed by Xenium spatial transcriptomics. Tumor regions (T) and histologically non-malignant pancreatic tissue (N) are indicated. (**B**) UMAP visualization of cell states identified by unsupervised Leiden clustering of the PDAC_10 Xenium dataset. Cluster 3 was annotated as invasive adenocarcinoma, whereas cluster 8 represented intraepithelial neoplasia. Clusters 5 and 6 corresponded to non-malignant pancreatic epithelial populations, comprising ductal/secretory epithelial cells and acinar-like epithelial cells, respectively. The remaining clusters represented fibroblast/CAF-like, T cell, myeloid, endothelial, endocrine/mast cell-like, and B/plasma-cell populations. (**C**) Dot plot showing representative epithelial marker-gene expression across Leiden clusters. Invasive adenocarcinoma (cluster 3) and intraepithelial neoplasia (cluster 8) displayed distinct epithelial transcriptional programs, whereas clusters 5 and 6 represented non-malignant ductal/secretory and acinar-like epithelial populations, respectively. The mutation-specific KRAS_G12V probe was excluded from the clustering procedure and mapped only after cluster annotation, providing independent genetic evidence for the identification of the neoplastic epithelial compartments. (**D**) Spatial distribution of the annotated epithelial compartments across the PDAC_10 Xenium tissue section. Invasive adenocarcinoma (cluster 3) and intraepithelial neoplasia (cluster 8) are shown separately from non-malignant pancreatic epithelial cells (clusters 5 + 6). Scale bar, 200 µm. (**E**) Spatial distribution of tumor-enriched clonotypes (ct5, ct2, ct4; T/N > 4) and non-enriched clonotypes (ct1, ct3, ct6; T/N < 1) on the PDAC_10 Xenium tissue section. Neoplastic epithelial cells (clusters 3 + 8) are shown in orange, selected clonotype T cells in red, and other cells in light grey. Tumor-enriched clonotypes preferentially localize within and around neoplastic epithelial compartments, whereas non-enriched clonotypes show a broader tissue distribution, as quantified in (F). The numbers of ct1– ct6 T cells were comparable between the T/N > 4 (n = 1,173) and T/N < 1 (n = 1,312) groups. Scale bar, 200 µm. (**F**) Distribution of single-cell distances of ct1–ct6 T cells to the nearest neoplastic epithelial cell in invasive adenocarcinoma (cluster 3), intraepithelial neoplasia (cluster 8), and the combined neoplastic epithelial compartment (clusters 3 + 8, distance to the nearest cell belonging to either cluster). Boxplots show the distribution of nearest-cell distances for each clonotype, with center lines indicating median distances. Clonotypes are ordered by decreasing tumor-to-normal (T/N) ratio, determined independently by bulk TCR repertoire sequencing; T/N ratios and numbers of Xenium-assigned T cells are indicated. Tumor-enriched clonotypes (ct5, ct2, ct4) showed shorter median distances to neoplastic epithelial cells than non-enriched clonotypes (ct1, ct3, ct6). (**G**) Fraction of marker-positive cells among ct1–ct6 T cells ordered by decreasing T/N ratio. The dashed line separates tumor-enriched from non-enriched clonotypes. Connecting lines are shown solely to visualize marker-expression patterns across clonotypes ordered by T/N ratio and do not represent longitudinal or temporal trajectories. GZMK, KLRG1, GZMA, and CD27 showed positive rank correlations with tumor enrichment (Spearman ρ = 1.00, 1.00, 0.94, and 1.00, respectively), whereas NKG7 showed a weaker correlation (ρ = 0.71). Together, these data link clonotype-level tumor enrichment to spatial proximity to neoplastic epithelial cells and a cytotoxic T-EM-related phenotype.

The spatial distribution of the clonotypes closely paralleled their independently determined T/N enrichment. Tumor-enriched clonotypes ct5, ct2, and ct4 preferentially localized within and around neoplastic epithelial compartments, whereas the non-enriched clonotypes ct1, ct3, and ct6 showed a broader tissue distribution (Fig. 8E). Quantitative single-cell distance analysis confirmed this pattern, with tumor-enriched clonotypes showing shorter median distances to neoplastic epithelial cells than non-enriched clonotypes across the analyzed compartments (Fig. 8F). Notably, ct5, which showed the strongest tumor enrichment, also displayed the shortest median distance to the combined neoplastic epithelial compartment. Thus, spatial proximity information, not contained in the bulk-derived T/N ratio, independently mirrored the clonotype-level pattern of tumor enrichment within the tissue architecture.

We next examined whether the effector-associated pattern observed by single-cell profiling was recapitulated in the spatial transcriptomics data. Across ct1–ct6, the fraction of GZMK-positive cells closely tracked T/N enrichment (Spearman ρ = 1.00, exact permutation *P* = 0.0028), with positive correlations for KLRG1 (ρ = 1.00, *P* = 0.0028), CD27 (ρ = 1.00, *P* = 0.0028), and GZMA (ρ = 0.94, *P* = 0.0167) (Fig. 8G). In contrast to single-cell profiling, NKG7 showed a weaker, non-significant correlation in the spatial analysis (ρ = 0.71, *P* = 0.136). Apart from this difference, these spatially resolved expression patterns were consistent with the T-EM-associated differentiation of tumor-enriched clonotypes observed independently by single-cell profiling.

Consistent with the findings for Cluster-A clonotypes in PDAC_49, the PDAC_10 analyses revealed concordant relationships between clonotype-level tumor enrichment, spatial proximity to neoplastic epithelial cells, and cytotoxic T-EM-associated differentiation.

The agreement between bulk TCR repertoire sequencing, single-cell transcriptional profiling, and Xenium spatial analysis supports spatial tumor engagement as an orthogonal feature. This spatial information may complement tumor-to-normal enrichment and transcriptional phenotype in prioritizing candidate tumor-reactive T cell clonotypes.

## Discussion

The identification of naturally occurring tumor-reactive TCRs remains a major challenge for the development of personalized TCR-based immunotherapies, particularly when the cognate tumor antigen is unknown and functional validation is limited by the availability of viable tumor material. Tumor-to-non-tumor (T/N) repertoire enrichment in combination with single-cell expression profiling provides an antigen-agnostic strategy to address this challenge by exploiting the selective expansion of T cell clonotypes within tumor tissue (9, 10).

Here, we extend this concept by showing that repertoire enrichment is linked to spatial tissue engagement and effector-associated transcriptional states within the tumor microenvironment. By integrating bulk and single-cell TCR profiling with clonotype-specific Xenium spatial transcriptomics, we demonstrate that tumor-enriched CD8⁺ T cell clonotypes preferentially associate with malignant tissue and display transcriptional features consistent with an antigen-experienced cytotoxic effector state. Thus, spatial analysis provides an orthogonal biological dimension that complements T/N repertoire enrichment for the identification and prioritization of candidate tumor-reactive TCRs (Fig. 9)

**Figure 9:**
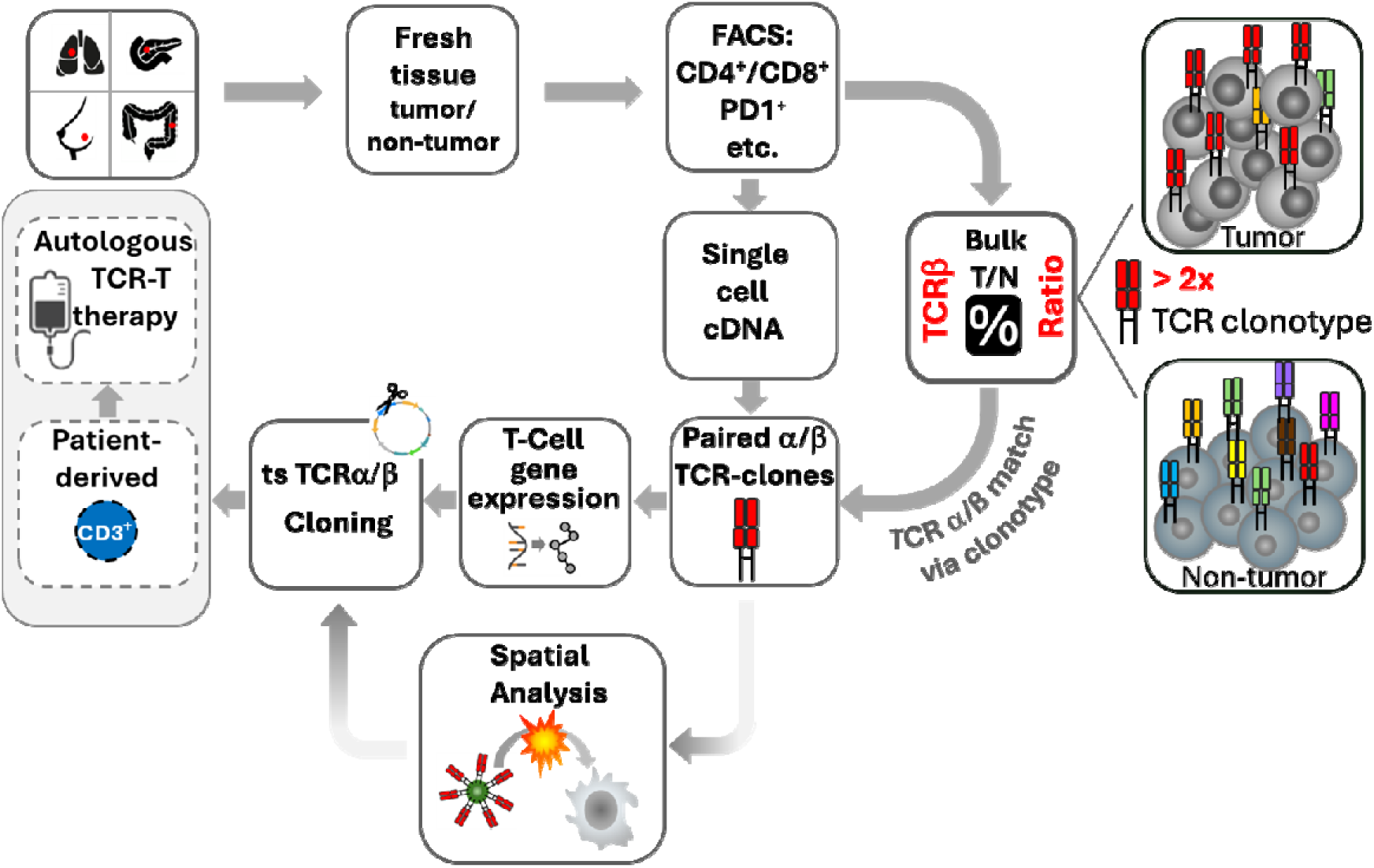
Antigen-agnostic identification and prioritization of candidate tumor-reactive T cell receptors (TCRs). Fresh tissue samples from solid tumors (NSCLC, CRC, BC, and PDAC) and matched adjacent non-tumor tissues are processed for analysis. Tumor-infiltrating T cells are isolated by FACS and subjected to single-cell analysis to pair TCR α and β chains and characterize their transcriptional phenotype. In parallel, bulk DNA from FACS-sorted T cells from tumor and matched non-tumor tissues is used to determine clonotype frequencies and calculate tumor-to-non-tumor (T/N) ratios. T cell clonotypes enriched at least twofold in tumor tissue are prioritized as candidate tumor-specific (ts) TCRs. Single-cell transcriptomic analysis provides complementary evidence of antigen-experienced effector and/or exhausted T cell states. As an orthogonal, antigen-agnostic approach to spatially support tumor reactivity, selected clonotypes can be mapped by Xenium spatial transcriptomics to assess their localization relative to malignant cells and tertiary lymphoid structures, with preferential tumor proximity providing additional evidence of tumor engagement. Prioritized TCRs may subsequently be selected for functional validation and development of autologous TCR-T cell therapy. Solid lines denote established experimental steps, whereas dashed lines indicate extensions toward therapeutic development.

This relationship was particularly evident for T cell Cluster-A, a population of highly similar HLA-A*02:01-restricted TCR clonotypes previously identified across multiple patients and solid tumor entities (13). In PDAC_49, Cluster-A CD8⁺ T cells were detected in tumor tissue but were absent from matched adjacent normal pancreas. Within the tumor, Cluster-A CD8⁺ T cells localized to malignant epithelial regions and were significantly closer to malignant cells than subcluster-defined naive-/T-CM-like CD8⁺ T cells. Their tumor-associated localization was accompanied by a predominantly cytotoxic effector-memory phenotype, providing independent spatial and transcriptional support for the tumor association inferred from T cell repertoire enrichment. Importantly, Cluster-A T cells were not restricted to a single anatomical niche, but were located both in proximity to malignant epithelial cells and within peri– and intratumoral tertiary lymphoid structures (TLS), indicating that tumor-associated clonotypes can occupy distinct spatial compartments within the same tumor microenvironment.

This association between T cell repertoire enrichment and spatial tumor engagement was not restricted to Cluster-A T cells. Analysis of the six most dominant CD8⁺ T cell clonotypes in PDAC_10 revealed a clear distinction between tumor-enriched and presumed bystander T cells. Clonotypes with progressively higher T/N ratios localized increasingly closer to malignant epithelial cells, showed higher expression of the effector-associated granzymes GZMA and GZMK, and preferentially mapped to T-EM clusters.

The concordance across independent clonotypes links three conceptually distinct measures, repertoire enrichment, spatial localization, and transcriptional phenotype, to tumor reactivity without having to know the cognate antigen. Rather than defining tumor reactivity by any single feature, our findings therefore support an integrated framework in which T/N enrichment provides the primary antigen-agnostic selection criterion, while spatial tumor engagement and effector-associated transcriptional states provide complementary evidence for prioritizing candidate tumor-reactive TCRs.

Transcriptional phenotypes of clonotypes analyzed across the two PDAC cases further support the described relationship between tumor-specific and bystander T cells. In PDAC_49, Cluster-A CD8⁺ T cells displayed a predominantly cytotoxic effector-memory (T-EM)-associated profile that was independently reproduced by single-cell RNA sequencing and Xenium spatial transcriptomics. Cluster-A cells showed increased representation of *GZMA*, *GZMK*, and *CCL5* compared with naive/central-memory-like CD8⁺ T cells, whereas *TCF7*, *CCR7*, and *SELL* were preferentially expressed by the latter population. Expression of *NKG7*, *PRF1*, *CXCR3*, *KLRG1*, and *CD27* in subsets of Cluster-A cells further indicated heterogeneity within an antigen-experienced effector-memory state rather than a uniformly terminally differentiated phenotype. A related GZMA/GZMK-associated pattern was observed among the tumor-enriched clonotypes in PDAC_10, suggesting that the association between tumor enrichment and an cytotoxic effector-memory-associated program is a shared feature of tumor-enriched T cell clonotypes in both samples.

One possible explanation for the antigen-experienced memory phenotype of Cluster-A T cells is recruitment of pre-existing memory T cells through microbial–tumor antigen cross-reactivity. Molecular mimicry between microbial and tumor-associated antigens is increasingly recognized as a potential contributor to anti-tumor T cell immunity, whereby T cells initially primed against microbial antigens can cross-recognize structurally related tumor-associated epitopes (16, 23). Such populations would already possess a memory phenotype and could rapidly respond upon encounter with a cross-reactive tumor antigen. This concept is particularly relevant as a few Cluster-A TCRs have previously been reported to recognize the EBV-derived LMP2A antigen as well as a potential tumor-associated target derived from TMEM161A (21). It is therefore plausible that Cluster-A T cells were initially primed during previous EBV infection and subsequently persisted as an antigen-experienced tissue-resident or central memory population (Supplementary Fig. S2). Importantly, EBER *in situ* hybridization demonstrated the absence of EBV-positive cells in the PDAC_49 sample, indicating that the pronounced intratumoral enrichment of Cluster-A was not driven by local EBV infection of B cells or epithelial cells. Instead, these findings are consistent with reactivation and expansion of a pre-existing cross-reactive memory population in response to a tumor-associated antigen. While TMEM161A has been proposed as a potential cross-reactive tumor target for related TCRs (21), the present study was not designed to establish the molecular identity of the recognized antigen, and our data therefore cannot confirm TMEM161A as the relevant tumor target. Similarly, the antigen specificities of the tumor-enriched PDAC_10 clonotypes remain unknown, and while microbial–tumor cross-reactivity may represent one possible explanation for their T-EM-associated phenotype, this possibility remains to be established.

Importantly, a high T/N ratio does not appear to define a single transcriptional state of tumor-reactive T cells. In our previous antigen-agnostic identification of KRAS Q61H-specific TCRs, clonotypes selected by strong tumor enrichment exhibited a more chronically activated and exhausted transcriptional program, including expression of *CXCL13*, cytotoxic effector genes, and multiple exhaustion-associated markers (10). Thus, both cytotoxic effector-memory-dominated and more chronically activated/exhausted T cell populations can be identified through tumor-selective repertoire enrichment. These observations suggest that T/N enrichment may capture antigen-driven accumulation within the tumor independently of the specific differentiation state of the responding T cells. Their phenotype may instead reflect their antigenic history and the nature and duration of antigen stimulation within the tumor.

The localization of Cluster-A cells within intratumoral TLSs adds a further dimension to this antigen-experienced phenotype. TLSs provide organized microenvironments for local antigen presentation and adaptive immune responses and have been associated with favorable clinical outcomes in PDAC (24). In the context of a potentially pre-existing antigen-experienced Cluster-A population, these structures may represent sites of local reactivation and expansion rather than sites of naive T cell priming. At the same time, Cluster-A cells were also found in close proximity to malignant epithelial cells, indicating that the same clonotype-defined population can occupy both lymphoid and tumor-engaged compartments within the tumor microenvironment. Although our spatial data cannot establish migration between these niches or identify the cells presenting the relevant antigen, this distribution is compatible with a model in which antigen-experienced Cluster-A cells are locally maintained or reactivated within TLSs and subsequently engage malignant tissue.

Beyond these biological insights, our findings have practical implications for antigen-agnostic TCR discovery. T/N enrichment can be determined from bulk TCR repertoire data and therefore provides a scalable primary criterion for identifying clonotypes that selectively accumulate within tumor tissue. Single-cell TCR/RNA sequencing can subsequently establish paired TCRαβ sequences and characterize their cellular phenotype, whereas clonotype-resolved spatial transcriptomics adds information on their localization relative to malignant and non-malignant tissue compartments. Spatial tumor association cannot establish cognate TCR–peptide–HLA recognition and therefore does not replace functional validation. However, particularly when viable tumor material or suitable autologous tumor models are unavailable, the convergence of repertoire enrichment, antigen-experienced cytotoxic effector phenotype, and spatial tumor engagement may provide a biologically informed basis for prioritizing candidate TCRs for subsequent functional testing. This becomes even more important when the Xenium analysis is extended to a multi-region spatial approach enabling selection of TCRs from clonotypes distributed in all regions of the tumor. Such TCRs may recognize antigens shared across spatially distinct tumor regions and could therefore provide a strategy for addressing intratumoral heterogeneity.

The potential relevance of this strategy extends beyond private tumor-reactive clonotypes: Cluster-A comprises highly similar TCRs identified across multiple HLA-A*02:01-positive patients and different solid tumor entities, suggesting convergent selection by a shared antigenic determinant. If their tumor-associated specificity and safety profile can be established, recurrent clonotype families such as Cluster-A may represent particularly attractive candidates for TCR-based therapies applicable across multiple patients.

## Limitations

The present spatial analyses were limited to two PDAC cases, restricting the generalizability of the observed relationships between T/N enrichment, spatial tumor association, and effector-associated transcriptional states. Although PDAC_49 and PDAC_10 provided complementary clonotype-level analyses, validation in larger cohorts and additional tumor entities will be required. Xenium provides targeted rather than transcriptome-wide expression profiling, and spatial proximity to malignant cells cannot establish cognate TCR–peptide–HLA recognition. Spatial evidence of tumor engagement should therefore be considered complementary to functional validation. Finally, the molecular tumor antigen recognized by Cluster-A remains unresolved, and the potential contribution of microbial– tumor cross-reactivity warrants further investigation.

## Conclusions

In conclusion, our findings extend tumor-to-non-tumor repertoire enrichment from a quantitative measure of intratumoral clonal expansion to an integrated framework for antigen-agnostic TCR discovery. Across two independent PDAC cases, tumor-enriched CD8⁺ T cell clonotypes preferentially associated with malignant tissue and displayed antigen-experienced cytotoxic effector phenotypes, while increasing T/N enrichment across independent clonotypes was accompanied by progressively closer tumor localization and increased GZMA/GZMK expression. These findings suggest that tumor reactivity may be inferred more robustly from the convergence of independent biological dimensions than from any single property of a T cell clonotype. In this framework, T/N enrichment provides the primary selection criterion, while transcriptional phenotype and spatial tumor engagement provide complementary evidence for prioritizing candidate tumor-reactive TCRs. Integration of these approaches may therefore help focus subsequent functional validation on clonotypes with the strongest biological evidence of tumor engagement and facilitate the development of antigen-agnostic strategies for therapeutic TCR discovery.

## Ethics statement

The study was performed in accordance with the declaration of Helsinki. Sample collection from patients was approved by the Ethics Committees of the Berlin Medical Association and Charité University Hospital Berlin (Eth-08/18, EA1/270/23, EA2/132/23) and signed informed consent was obtained from all patients.

## Data and code availability statement

The custom analysis scripts developed for this study will be made publicly available through GitHub, and derived data underlying the figures and quantitative analyses will be deposited in appropriate public repositories upon publication of the article. Patient-derived raw data that cannot be made publicly available due to ethical, legal, or privacy restrictions will be made available to qualified researchers upon reasonable request and in accordance with applicable data protection and consent requirements.

## Author contributions

VS: Conceptualization, Methodology, Software, Validation, Formal analysis, Investigation, Data curation, Visualization, Project administration, Supervision, Funding acquisition, Writing – original draft, Writing – review & editing. JF: Conceptualization, Software, Validation, Formal analysis, Data curation, Visualization, Writing – original draft, Writing – review & editing. KG: Conceptualization, Software, Validation, Formal analysis, Data curation, Visualization, Writing – original draft, Writing – review & editing. BH: Conceptualization, Methodology, Validation, Investigation, Resources, Data curation, Visualization, Writing – original draft, Writing – review & editing. SS: Methodology, Validation, Investigation, Resources, Data curation, Writing – review & editing. AD: Methodology, Validation, Investigation, Data curation, Writing – review & editing. MD: Methodology, Validation, Investigation, Resources, Data curation, Writing – review & editing. NG: Methodology, Validation, Investigation, Resources, Data curation, Writing – review & editing. SP: Methodology, Validation, Investigation, Resources, Data curation, Writing – review & editing. DB: Software, Validation, Formal analysis, Data curation, Writing – review & editing. JG: Methodology, Validation, Formal analysis, Data curation, Writing – review & editing. TC: Methodology, Validation, Formal analysis, Investigation, Data curation, Writing – review & editing. CL: Methodology, Validation, Investigation, Data curation, Writing – review & editing. AT: Methodology, Investigation, Writing – review & editing. MM: Validation, Resources, Writing – review & editing. DH: Validation, Resources, Writing – review & editing. TK: Validation, Formal analysis, Writing – review & editing. DK: Validation, Resources, Writing – review & editing. KS: Resources, Data curation, Writing – review & editing. MH: Formal analysis, Resources, Visualization, Writing – review & editing. SH: Conceptualization, Methodology, Software, Formal analysis, Resources, Project administration, Funding acquisition, Writing – original draft, Writing – review & editing. VL: Conceptualization, Methodology, Validation, Formal analysis, Data curation, Visualization, Project administration, Funding acquisition, Writing – original draft, Writing – review & editing. SE: Conceptualization, Methodology, Validation, Investigation, Resources, Data curation, Visualization, Project administration, Supervision, Funding acquisition, Writing – original draft, Writing – review & editing.

## Generative AI statement

Generative artificial intelligence (OpenAI ChatGPT) was used during manuscript preparation to assist with English-language editing, improve clarity and readability, and support the development, refinement, and troubleshooting of computational code. Generative AI was not used to generate experimental data or to independently perform scientific interpretation or draw conclusions. All AI-assisted text and code were critically reviewed, verified, and, where necessary, modified by the authors. The authors retain full responsibility for the scientific content, accuracy, and integrity of the manuscript and all reported analyses.

## Acknowledgments

We are grateful to Prof. Dr. Rudolf Hammer for helpful comments and discussions concerning the manuscript. We would like to express our gratitude to the Charité 3^R^ Primary Tissue Pipeline and clinical teams for their assistance with tissue collection. We also thank the EU and Investitionsbank Berlin (IBB) for supporting parts of the work on colorectal, breast, and pancreatic cancer through the EFRE and ProFIT funding instruments.

