## Supplementary Information for "Tumor-versus-non-tumor T cell clonotype enrichment enables antigen-agnostic prioritization of tumor-reactive TCR candidates, supported by single-cell and spatial phenotyping"

**Supplementary Results and Figures**

**Independent reanalysis of published NSCLC TCR repertoire data supports tumor enrichment of Cluster-A-associated TCRβ clonotypes**

To independently assess the tissue distribution of Cluster-A clonotypes in an external cohort, we reanalyzed previously published TCRβ repertoire data from patients with non-small cell lung cancer (NSCLC), comprising tumor tissue, matched adjacent non-tumor lung tissue, and peripheral blood (1). Specifically, we examined the clonotype-level data reported in Fig. 2F of Chiao et al., focusing on TCRβ CDR3 sequences compatible with the Cluster-A family based on CDR3 sequence homology and TRBV10-2 usage (1).

The identified Cluster-A-associated CDR3β sequences showed variable distributions between tumor and matched adjacent lung tissue (Table S1). Six of the eight sequences showed higher mean clonal frequencies in tumor than in adjacent lung, whereas one sequence was detected only in adjacent lung and peripheral blood and one showed a slightly higher mean frequency in adjacent lung than in tumor (Table S1). All eight sequences were detectable in peripheral blood, generally at substantially lower frequencies than in tissue. Overall, these independent data support a preferential tumor-associated distribution of Cluster-A-related TCRβ sequences, while also demonstrating their low-frequency detection in peripheral blood.

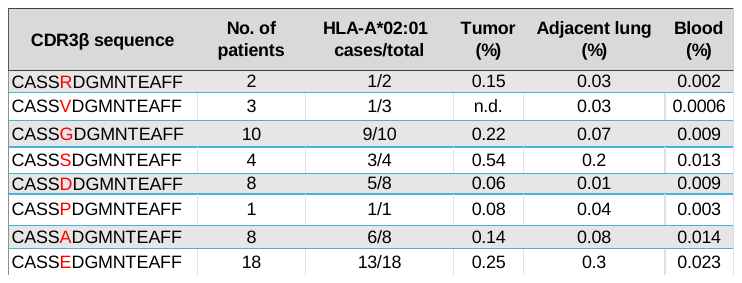

**Table S1:**

**Distribution of Cluster-A-associated TRBV10-2 CDR3β sequences in a published NSCLC cohort.** Cluster-A-associated CDR3β sequences were identified from the clonotype-level data reported in Fig. 2F of Chiao et al. based on CDR3β sequence homology and TRBV10-2 usage (1). For each sequence, the number of patients in whom it was detected, the number of HLA-A*02:01-positive cases relative to the total number of cases, and the reported mean clonal frequencies (% per patient) in tumor tissue, matched adjacent non-tumor lung tissue, and peripheral blood are shown. Values represent data reported by Chiao et al. (1). n.d., not detected.

**EBV-associated cells were absent in PDAC_49**

Because a small subgroup of Cluster-A TCRs have previously been reported to recognize the EBV antigen LMP2A, we asked whether local EBV infection could account for the accumulation of Cluster-A cells in PDAC_49. To address this, we performed EBER *in situ* hybridization using a colorectal carcinoma (CRC) case with EBER-positive cells as a positive assay control.

EBER *in situ* hybridization detected no EBER-positive cells in PDAC_49, including malignant epithelial cells and the corresponding normal pancreatic ductal epithelium, whereas only rare EBER-positive cells were observed in the CRC control tissue (Fig. S1A, B).

These findings argue against ongoing local EBV infection as the primary driver of Cluster-A accumulation in PDAC_49 and are compatible with a model in which pre-existing Cluster-A T cells expand locally in response to cross-reactive tumor-associated antigens.

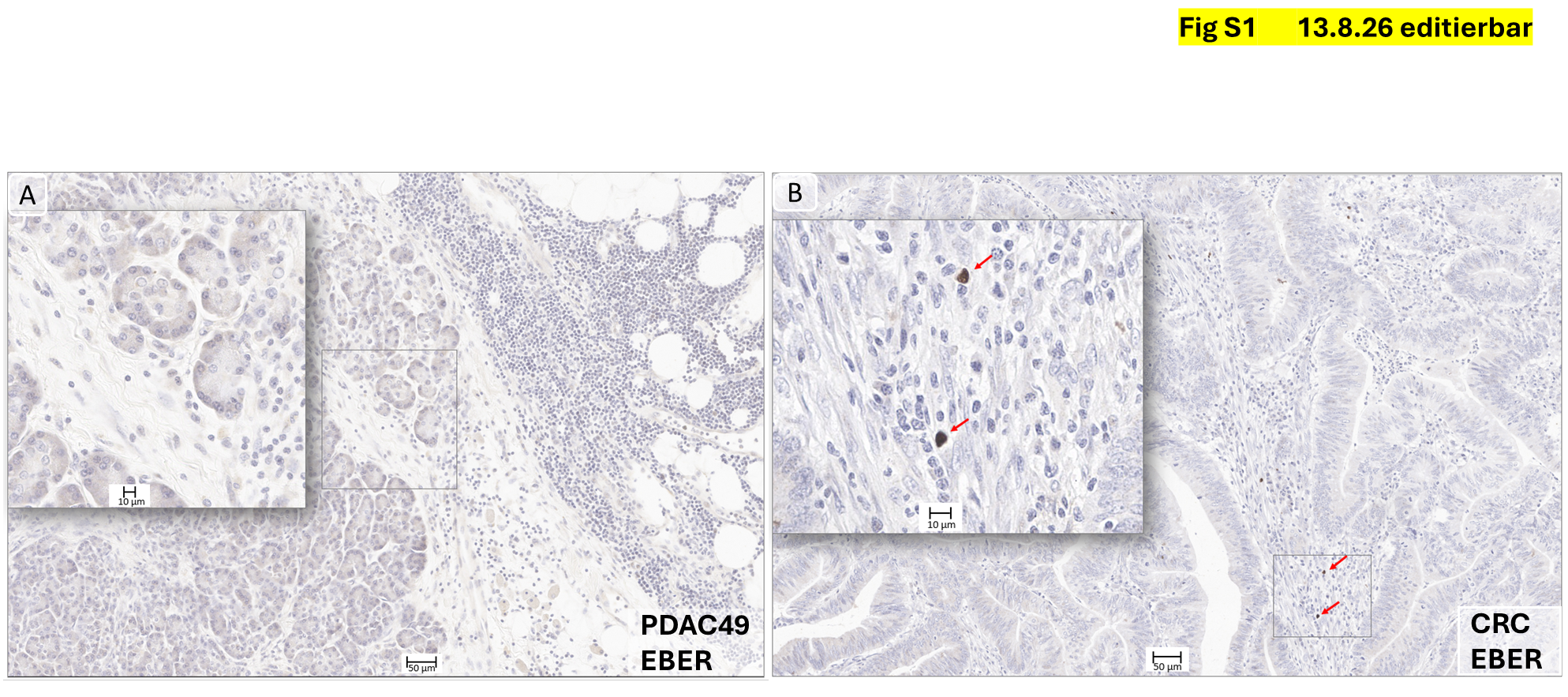

**Figure S1:**

**EBER *in situ* hybridization demonstrates the absence of EBV-infected cells in PDAC_49. (A)** No EBER-positive cells were detected in PDAC_49, including malignant epithelial cells and the corresponding normal pancreatic ductal epithelium. These findings argue against ongoing local EBV infection as the driver of Cluster-A T cell accumulation in PDAC_49. **(B)** Colorectal carcinoma (CRC) positive control showing rare EBER-positive cells (arrows), consistent with occasional EBV-infected lymphoid cells.

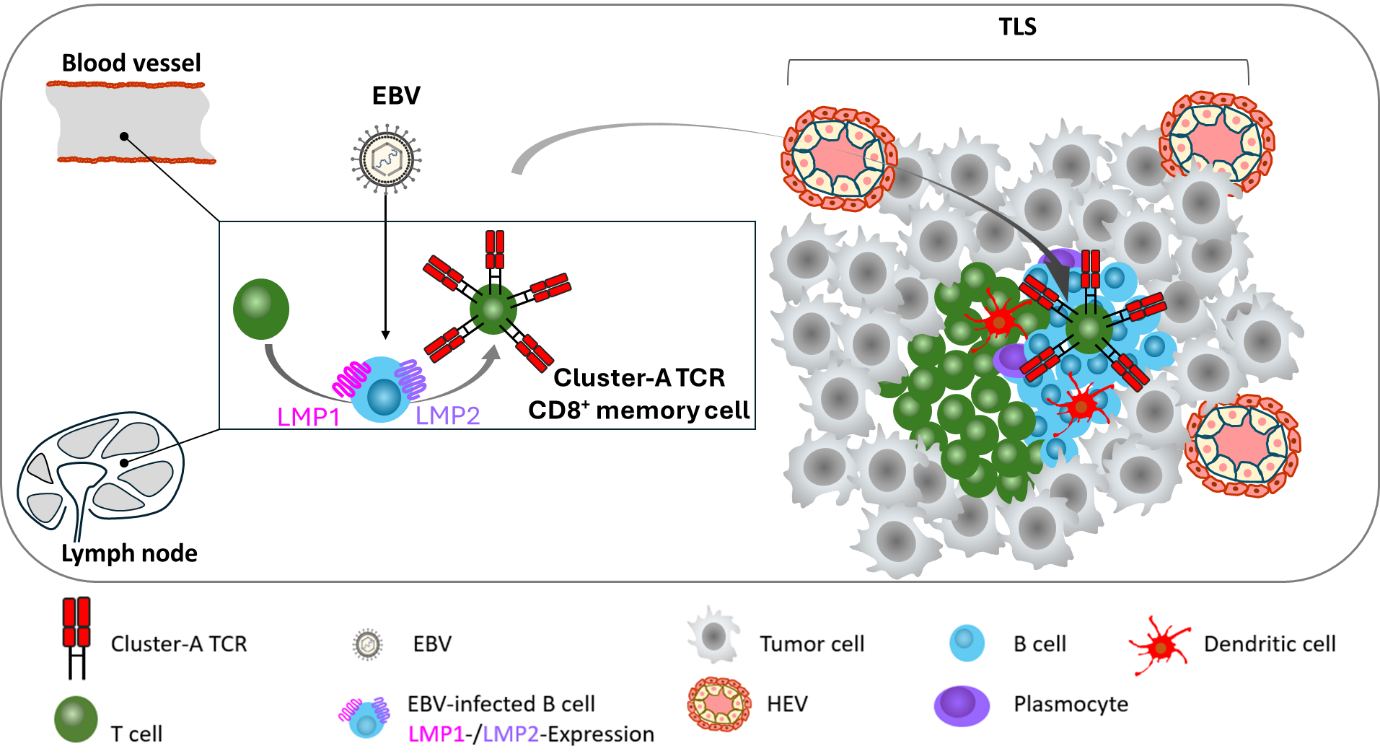

**Figure S2:**

**Proposed model of prior EBV priming and subsequent tumor-associated expansion of cross-reactive Cluster-A CD8⁺ T cells.** Proposed initial priming of Cluster-A CD8⁺ T cells during a previous EBV infection. EBV-infected B cells expressing the viral antigens LMP1 and LMP2 may activate T cells bearing recurrent Cluster-A TCRs, resulting in the establishment of circulating memory T-cell populations. The detection of Cluster-A clonotypes at low frequencies in peripheral blood is compatible with such a systemic memory reservoir but does not establish an EBV origin. Within the tumor microenvironment, pre-existing Cluster-A memory T cells may undergo preferential recruitment and/or local re-expansion following recognition of cross-reactive tumor-associated antigens. Their enrichment in tumor tissue, localization in close proximity to malignant cells and within tertiary lymphoid structures (TLS), and predominantly antigen-experienced effector phenotype are consistent with local tumor engagement. The absence or extreme rarity of EBV-positive cells in the analyzed tumor tissue argues against ongoing local EBV infection as the primary driver of Cluster-A T cell expansion. Together, these observations are compatible with a model in which prior EBV exposure generates cross-reactive memory T cells that subsequently expand in response to tumor-associated antigens.
